# Rapid divergence in sperm morphology and transgressive segregation of hybrid sperm kinematics in a young pupfish radiation

**DOI:** 10.64898/2026.08.17.745335

**Authors:** Oskar Golwala, Christopher H. Martin, Matthew C. Kustra

## Abstract

Understanding how divergence in reproductive traits can promote speciation remains a fundamental question in evolutionary biology. Sperm morphology and kinematics diverge rapidly across species. However, the effect of hybridization between recently diverged species on sperm traits remains unclear, limiting our understanding of how reproductive isolation evolves. Here, we evaluated sperm morphology, kinematics, and trait integration in species of a young (∼10,000 years), sympatric *Cyprinodon* pupfish radiation from San Salvador Island, Bahamas, as well as fertile advanced-generation hybrids between two of these species. We found significant divergence in flagellum length, midpiece area, and sperm velocity among species. In contrast, hybrids displayed transgressive kinematic profiles defined by high velocities, reduced path curvature, and distinct patterns of sperm kinematic integration. Our findings suggest that hybridization between recently diverged species may reorganize the underlying control of sperm locomotor mechanisms, generating novel phenotypes that could contribute to reproductive isolation in the early stages of speciation.

## Introduction

Identifying the reproductive isolation mechanisms that maintain species boundaries in young adaptive radiations is crucial for understanding speciation (El Nagar & MacColl, 2016; Fitzpatrick, Montgomerie, et al., 2009; Lackey & Boughman, 2017; Martin & Richards, 2019; Rometsch et al., 2020). Analyzing reproductive traits is especially informative as they rapidly evolve and are often of key importance in sympatric speciation where geography does not contribute to reproductive isolation (Gillespie et al., 2020; Kagawa et al., 2023; Lifjeld, Garcia-del-Rey, et al., 2025; Wagner et al., 2012).

Sperm have an incredible amount of morphological diversity across taxa (Fitzpatrick et al., 2020; Pitnick et al., 2009; Torgerson et al., 2002). Sperm divergence is driven by strong, species-specific selective forces (e.g., fertilization environment, female-male interactions) (Fitzpatrick & Lüpold, 2014; Kahrl et al., 2021; Syed et al., 2025), that can reinforce different sperm optima (Kleven et al., 2008). Species differences in sperm may therefore play an important role in the evolution of reproductive incompatibilities (i.e., postmating prezygotic reproductive barriers) (Garlovsky et al., 2024; Kustra et al., 2025; Lifjeld, Cramer, et al., 2025; Westram et al., 2022) as well as hybrid sterility (i.e., postzygotic reproductive barrier) (Coyne & Orr, 1997). However, little is known about whether fertile hybrids between recently diverged species exhibit weak reproductive incompatibilities (Schumer et al., 2015), further limiting our understanding of the initial reproductive isolating mechanisms contributing to speciation.

Sperm may offer further insight into genetic incompatibilities within hybrids because of their high level of biomechanical integration and functional complexity (Gaffney et al., 2011; Pereira et al., 2017). Biomechanically complex systems have come to the forefront of speciation research due to their capacity for transgressive performance (Holzman & Hulsey, 2017; Parnell et al., 2008; Rieseberg et al., 1999). These novel hybrid phenotypes are the result of new morphological trait combinations that can result in unique performance profiles (Bell & Travis, 2005; Evans & Felice, 2026; Holzman & Hulsey, 2017; Parnell et al., 2008; Rieseberg et al., 1999). Although sperm share many similarities with previously studied biomechanical systems where morphology, energetics, and sensory systems all modulate performance (Fitzpatrick, Craig, et al., 2009; Fitzpatrick et al., 2020; Fitzpatrick, Montgomerie, et al., 2009; Fitzpatrick & Lüpold, 2014; Gallego et al., 2013; Humphries et al., 2008; Safran et al., 2013), investigation of sperm morphology and transgressive kinematics in hybrids remains lacking.

The San Salvador Island pupfish adaptive radiation consists of four species: a generalist (*Cyprinodon variegatus*) and three trophic specialists: a molluscivore (*C. brontotheroides*), a scale-eating specialist (*C. desquamator*), and an undescribed intermediate scale-eater (*C.* sp. ‘wide-mouth’) (Martin & Wainwright, 2011, 2013a, 2013b) (Fig. 1A). Within only 10 ky, these species diverged rapidly in craniofacial morphology and behavior (Martin & Wainwright, 2013a; Palominos et al., 2023, 2025; Patton et al., 2022; Richards et al., 2021; Richards & Martin, 2017). Hybrids among these species are fertile and viable in the laboratory, but have reduced density- and frequency-dependent rates of growth and survival in field enclosures on San Salvador Island (Martin, 2016; Martin & Gould, 2020; Martin & Wainwright, 2013b) and show gene misexpression during early development (McGirr & Martin, 2019, 2021). Because pupfishes are external fertilizers with competitive mating systems (Kodric-Brown, 1977; Leiser & Itzkowitz, 2004), postcopulatory sexual selection and potential hybrid gametic dysfunction may serve as critical mechanisms maintaining species boundaries when prezygotic barriers are weak (Farnitano & Sweigart, 2023; Presgraves, 2010; Price, 1997). This young system, therefore, provides a great opportunity to investigate divergence in sperm morphology and kinematics within a recent adaptive radiation and test whether hybridization can reshape sperm morphology and swimming mechanics.

**Fig. 1:**
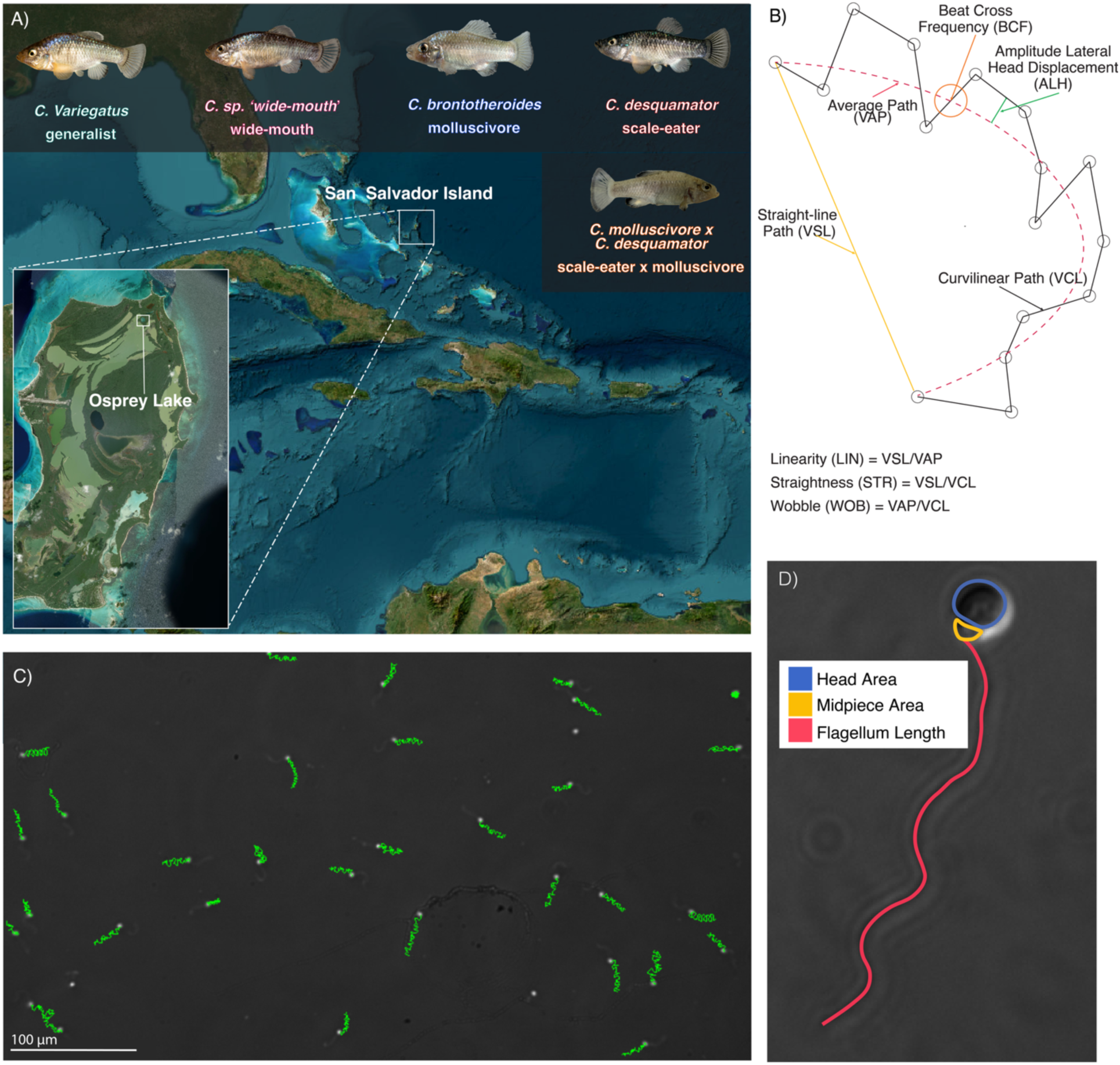
Sperm morphology and kinematic measurements. (A) Map of San Salvador Island in the Bahamas along with the pupfish species occupying Osprey Lake and lab-reared scale-eater x molluscivore hybrids. (B) Cartoon depictions of measured and derived kinematic variables. True kinematic variables incorporate time to measure rates and velocities. (C) Sample frame from a measured molluscivore sperm sample. Green lines show tracked path for motile sperm. (D) Sperm morphology measurements overlaid on a brightfield image.

Here, we examined how sperm morphology and kinematics differ among lab-reared pupfish species and advanced-generation hybrids between the molluscivore and scale-eater. We first quantify how sperm morphology and velocity have diverged among species. We then test for transgressive segregation in hybrid sperm and use underlying trait variance-covariance matrices to estimate how hybridization reshapes functional correlations.

## Materials and Methods

### Fish Collection and Husbandry

We used seine nets to sample Osprey Lake, San Salvador Island, Bahamas, for four species of pupfish: a generalist (*C. variegatus*), a molluscivore (C*. brontotheroides*), a scale-eater (*C. desquamator*), and an undescribed *C.* sp. ‘wide-mouth’ in July of 2017 and March of 2018. Because these species form a closely related genetic cluster, phylogenetic correction was unnecessary. This study used a combination of fifth-, sixth-, and seventh-generation pupfishes derived from this collection as well as an F2 and F3 hybrid intercross population reared from a single molluscivore (*C. brontotheroides*) and scale-eater (*C. desquamator*) pair from Osprey Lake. Fish were reared in a flow-through setup to maintain consistency across tanks. All animal procedures used in this study were approved by the UC Berkeley Animal Care and Use Committee (AUP-2021-07-14515, AUP-2021-02-14062-1, AUP-2020-02-12986, AUP-2018-08-11373).

We collected unique sperm samples from each of 59 individuals. Six samples were discarded because they lacked sufficient (<15) sperm for accurate morphology measurements and one sample was discarded because after filtering, the remaining sample size for kinematics was below 100. Therefore, our results consist of data from 12 generalists, 10 wide-mouths, 10 molluscivores, 10 scale-eaters, and 10 F2 and F3 hybrids.

### Sperm collection

We first anesthetized male pupfish using a dilute concentration of tricaine methanesulfonate (MS-222) and stripped milt using manual palpation into 1 µL microcapillary tubes to capture the full milt sample from each fish. We mixed the milt with 4 µL of Hank’s buffer solution (HBSS; Life Technologies, USA) in 1.5 mL microcentrifuge tubes to dilute sperm for accurate kinematic measurements while also extending motility before sample activation (Jing et al., 2009; Sayyari et al., 2024). We then used 1 µL of the HBSS-milt sample for kinematic analysis and the remaining volume was fixed and plated for morphological measurement.

### Sperm motility

We activated sperm by diluting 1 µL of the HBSS-milt mixture into a separate microcentrifuge tube with 3 µL of fresh tank water. These volumes were chosen to optimize concentration for kinematic data collection. We then added 3 µL of activated milt into a four-chambered 3 µL Leja slide (Leja Products B.V., Netherlands). We recorded individual sperm motility at 50 frames per second with a Nikon Ci-L microscope with a 10x phase-contrast objective, as recommended by Microptic computer-aided sperm analysis (CASA) (Microptic, S.L., Spain). Sperm motility was analyzed with Microptic CASA software for 2 minutes and 30 seconds. We moved the field of view after each captured field to record as many unique sperm as possible. Post-capture, we excluded clumped sperm and fields with obvious disturbances that led to faulty kinematic profiles. To ensure higher accuracy in estimated sperm motility measurements, we additionally excluded any sperm that had less than the maximum frames tracked (<49).

Raw kinematic data consisted of 8 variables: curvilinear velocity (VCL), straight-line velocity (VSL), average path velocity (VAP), linearity (LIN), straightness (STR), wobble (WOB), amplitude of lateral head displacement (ALH), and beat cross frequency (BCF) (Fig. 1B). To reduce the influence of immotile and low-quality sperm on average kinematic estimates, we excluded sperm with very low curvilinear velocity using a conservative threshold (VCL > 15) following methods from Kustra et al. (2026). This threshold corresponded to the valley of the bimodal VCL distribution, to which the left peak represents immotile or minimally motile sperm (Fig. S1).

### Sperm morphology

We mixed the remaining HBSS-milt mixture with 8 µL of 4% paraformaldehyde (PFA) diluted with phosphate-buffered saline (PBS). Milt samples were left to fix for 30-60 minutes as this maximized sperm integrity. We then spread samples across charged microscope slides and allowed them to airdry before rinsing with deionized (DI) water.

We imaged sperm samples using 100x brightfield illumination on a Zeiss AxioImager M2 microscope using the QIClick digital CCD to capture high-bit depth gray scale images at 100x magnification. We measured flagellum length, total length, head area, and midpiece area manually using FIJI (ImageJ) (Fig. 1D). Head-tail ratio was later calculated to complement existing measurements because of its relevance for biomechanical efficiency (Hook & Fisher, 2020; Humphries et al., 2008). We later converted measurements from pixel values to µm for lengths and µm^2^ for area measurements. We measured between 16 and 30 sperm in each milt sample; for feasibility, we reduced our sample size to ∼20 from ∼30 after preliminary testing showed that the reduction in error between 20 and 30 sperm was marginal (Fig. S2).

### Statistical Analyses: Divergence in sperm morphology and velocity among species

To test for species differences in sperm morphology and sperm velocity while accounting for repeated sperm measurements within males, we used linear mixed-effects models from the lme4 package with species as a fixed effect and male identity as a random effect (Bates et al., 2015). The significance of species effects was assessed using one-way ANOVAs with a Tukey’s pairwise post-hoc test between species (emmeans package; Lenth & Piaskowski, 2026). For sperm velocity, we performed a PCA on VSL, VCL, and VAP values on all sperm and used the first principal component as an overall measure of sperm velocity (Fig. S3). These velocity variables were included as each has been associated with fertilization performance in fishes (Gallego et al., 2013; Gallego & Asturiano, 2019; Gasparini et al., 2010; Kurta et al., 2022). To see if any differences in velocity were the result of differences in morphology, we also conducted linear models for each morphology-velocity combination (see supplemental methods).

We additionally quantified the intra-individual coefficient of variation (CV) for each morphological trait and used linear model mean adjusted standard deviation (SD_adj_) for composite velocity to measure variance. We used CV due to established relationships between intra-individual morphology coefficient of variation (CV) and strength of selection for a given trait (Calhim et al., 2007; Immler et al., 2011; Kleven et al., 2008) and used SD_adj_ for velocity as it both accounts for mean-variance scaling in accordance with Taylor’s power law (Taylor, 1961) and accommodates near-zero mean values like those produced in our velocity PCA (Houle, 1992). We calculated CV using the standard deviation of natural log-transformed morphological traits (SD_ln_). Log-transformation eliminated mean-scale dependencies across all morphological traits (Fig. S4; Hosken et al., 2025; Kleven et al., 2008). We used one-way ANOVAs with a Tukey’s pairwise post-hoc test between species to test for significant differences in mean intra-individual CV.

We checked to see if any differences in kinematics were the result of differences in morphology by conducting linear models for each morphology-velocity combination (head-area to flagellum-length ratio was included as a morphology trait in this analysis due to its demonstrated relationship with velocity; Humphries et al., 2008)

### Statistical Analyses: F2 hybrid sperm transgression and integration

We analyzed differences in hybrid sperm distributions by mapping density distributions for each morphological and kinematic variable. For each individual variable combination, we calculated the first three statistical moments of the distribution: mean, variance, and skewness (moments package; Komsta & Novomestky, 2022). We calculated standardized effect sizes (Cohen’s *d*) and 95% confidence intervals for hybrids relative to the pooled parental species average (molluscivores and scale-eaters; Cohen, 1988). To compare hybrids and parents on a single, shared scale, we mapped both parental species means onto this same coordinate system as standard deviation offsets from the parental baseline (zero). We defined a trait moment as transgressive when both parental means fell entirely outside the hybrid’s 95% confidence interval.

We quantified sperm morphological and kinematic integration for each species to identify how phenotypic changes could lead to unique hybrid sperm phenotypes (Moore et al., 2013; Pigliucci, 2003). We calculated correlation matrices for both morphological and kinematic traits at the individual level and then took means to get species-level correlation matrices. We used relative eigenvalue variance (*V_rel_)* as a measure of concentration of integration, quantifying how much intraspecific variation is channeled along a single multivariate axis (Chan et al., 2024; Conaway & Adams, 2022; Pavlicev et al., 2009). We used ANOVAs and Tukey’s pairwise comparisons to test whether hybrids displayed significantly elevated or depressed *V_rel_*.

To evaluate whether species share conserved integration architectures, we compared correlation structures across species using multivariate permutation tests and matrix subspace comparisons (Machado et al., 2018; Marroig & Cheverud, 2001; Porto et al., 2009). First, we calculated Fisher *z*-transformed individual correlation matrices and tested for overall species differences using a permutation-based MANOVA (Adams & Collyer, 2018; Anderson, 2008). To confirm that results reflected true structural divergence rather than variance artifacts, we verified multivariate dispersion homogeneity (Anderson, 2006) before running post-hoc pairwise Bonferroni corrected permutation MANOVAs on species pairs. Second, to quantify functional matrix alignment, we calculated Krzanowski distances across leading principal components, directly measuring subspace overlap between species correlation structures (Aguirre et al., 2014; Horta-Lacueva et al., 2021; Krzanowski, 1979; Penna et al., 2017; Rohner, 2025)

## Results

### Sperm morphology and velocity have rapidly diverged

We found significant differences in flagellum length (one-way ANOVA, *F*_3,39_ = 4.43, *P* = 0.009) and midpiece area (*F*_3,40_ = 9.26, *P* < 0.001) among non-hybrid species in the radiation. In contrast, head area did not differ significantly among species (*F*_3,38_ = 2.24, *P* = 0.100). Post-hoc estimated marginal means comparisons further elicited which species pairs differed significantly for each trait (Fig. 2A-C). Wide-mouths and scale-eaters both exhibited significantly higher midpiece areas than the generalists (wide-mouth: -0.20 ± 0.07, *P* = 0.039; scale-eater: -0.20 ± 0.07, *P* = 0.033) and the molluscivores (wide-mouth: -0.30 ± 0.07, *P* < 0.001; scale-eater: -0.30 ± 0.07, *P* < 0.001) (Fig. 2B; Table S1). Additionally, wide-mouths showed significantly shorter flagellum lengths than the scale-eaters (-3.31 ± 0.88, *P* = 0.004) (Fig. 2C; Table S1). Pairwise comparisons focused on the hybrids revealed that they only differed from wide-mouths in flagellum length (3.04 ± 0.90, *P* = 0.012) (Fig. 2C; Table S1), showing similar lengths to both scale-eaters and generalists.

**Fig. 2:**
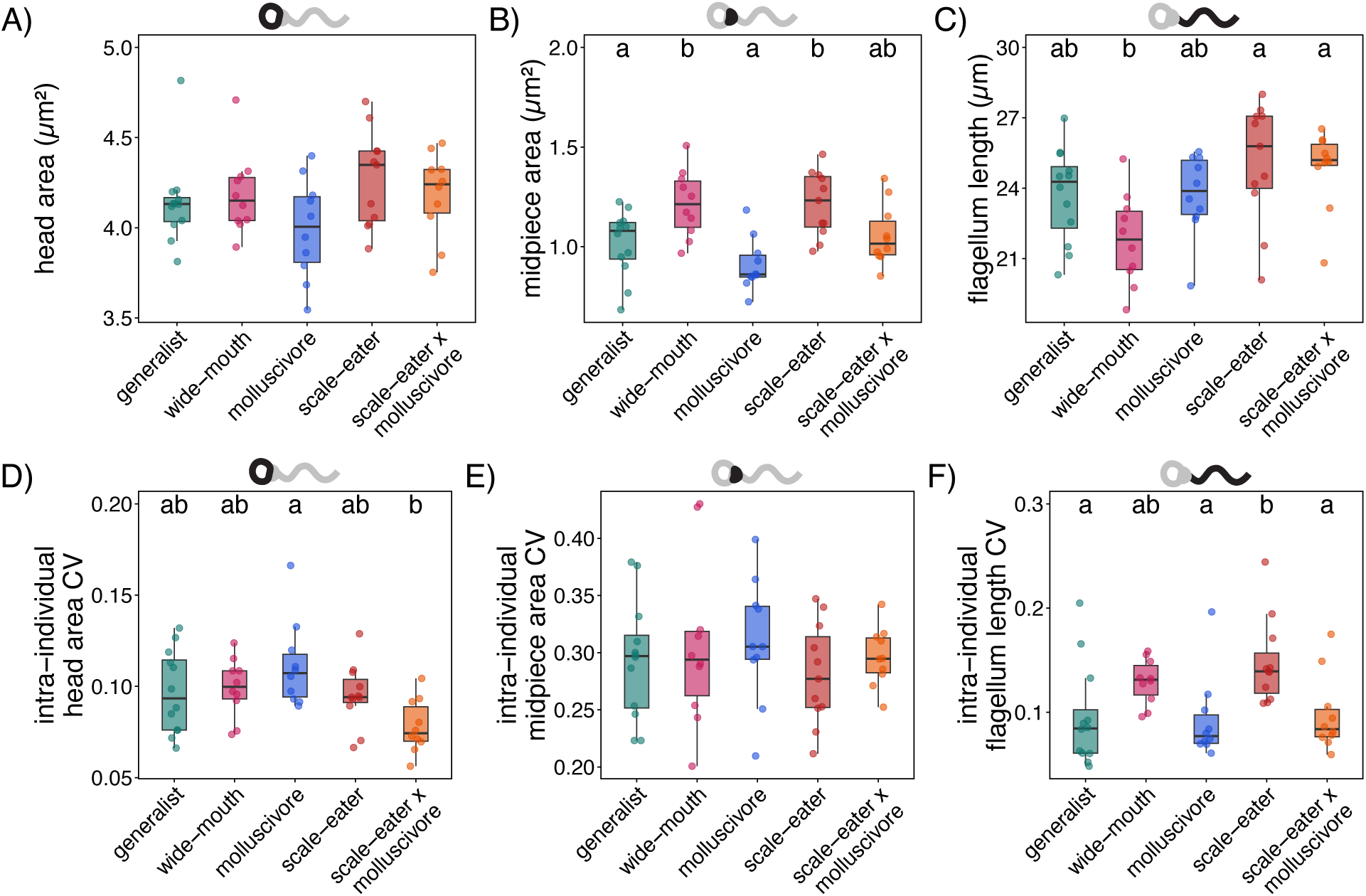
Rapid divergence in sperm morphology among species. (A) Mean head area, (B) midpiece area, and (C) flagellum length across species using male averages of all sperm morphology measurements. Intra-individual coefficient of variation (CV) of (D) head area, (E) midpiece area, and (F) flagellum length across species. Points are the average value of individual males. Species that do not share a letter grouping significantly differed in Tukey’s post hoc pairwise comparisons.

Species also differed in intra-individual morphological coefficient of variation (CV). Head area CV differed significantly among species (*F*_4,48_ = 3.85, *P* = 0.009), as did flagellum length CV (*F*_4,48_ = 4.29, *P* = 0.005). Midpiece CV did not differ significantly between species (*F*_4,48_ = 0.520, *P* = 0.722). Pairwise comparisons revealed that molluscivores showed higher variation than hybrids in head area (0.03 ± 0.01, *P* = 0.003), while scale-eaters had higher flagellum length CV than generalists (0.05 ± 0.02, *P* = 0.018), molluscivores (0.06 ± 0.02, *P* = 0.019), and hybrids (0.05 ± 0.02, *P* = 0.040) (Fig. 2D-F; Table S1).

Our integrated velocity metric, the first principal component of sperm velocity metric PCA (PC1), accounted for 83% of the total variation across the three velocity traits and loaded positively with all three traits (VSL = 0.55, VCL = 0.57, VAP = 0.61; Fig. S3). Among non-hybrid species, we found significant differences in velocity (*F*_3,43_ = 3.70, *P* = 0.019; Fig. 3A), with molluscivores having a higher mean composite velocity than scale-eaters (0.92 ± 0.31, *P* = 0.037) (Table S2).

**Fig. 3:**
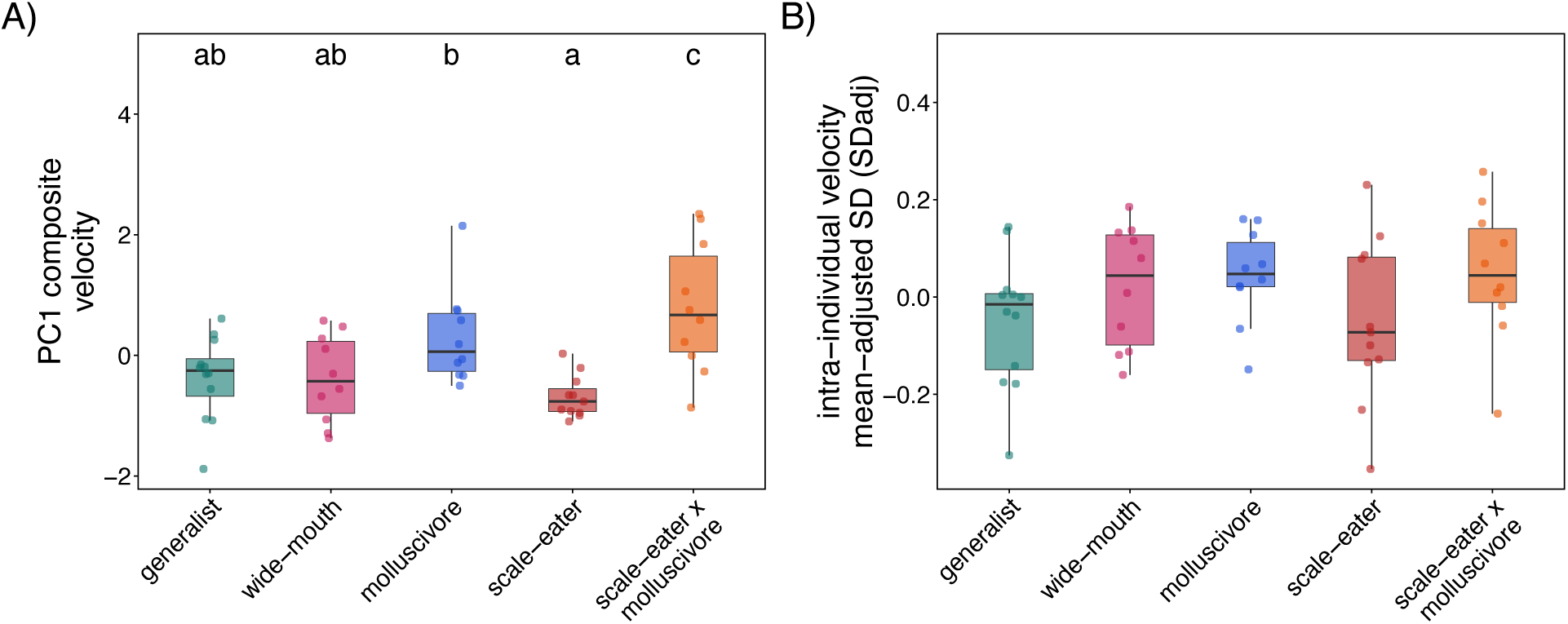
Sperm velocity, but not variance, substantially increased in hybrids. (A) Mean composite sperm velocity for each species based on averages. (B) Intra-individual coefficient of variation for sperm velocity across species. Points are the average value of individual males, while distributions for individual sperm are shown on the right of each boxplot.

Hybrids showed velocity metrics that were significantly higher than all species (Fig. 3A). This increase in velocity was not the result of decreased longevity (Fig. S5) or one component of the PCA outweighing others (Fig. 3, S3). The intra-individual velocity adjusted standard deviation (SD_adj_) showed no species effects (*F*_4,48_ = 1.41, *P* = 0.246; Fig. 3B). Species also showed no significant differences in morphology-kinematic relationships indicating that the demonstrated species differences in morphology do not have divergent impacts on kinematic scaling (see supplemental results; Fig. S6).

### Transgressive kinematics in hybrid sperm

Our analyses of phenotypic distributions in kinematic traits revealed pervasive phenotypic transgression while morphological variables failed to show similar distribution differences (Fig. 4; Fig. S7). Analyses of distribution means demonstrated that hybridization created high-velocity hyper-linear sperm that significantly exceeded the parental baseline. This was marked by transgressive means in straight-line velocity (VSL: *d* = 1.942, CI = [1.05,2.80]), linearity (LIN; *d* = 2.298, CI = [1.36, 3.21]), straightness (STR; *d* = 2.201, CI = [1.28, 3.10]), beat-cross frequency (BCF; *d* = 1.484, CI = [0.65, 2.30]), and wobble (WOB; *d* = 1.426, CI = [0.60, 2.23]) (Fig. 4).

**Fig. 4:**
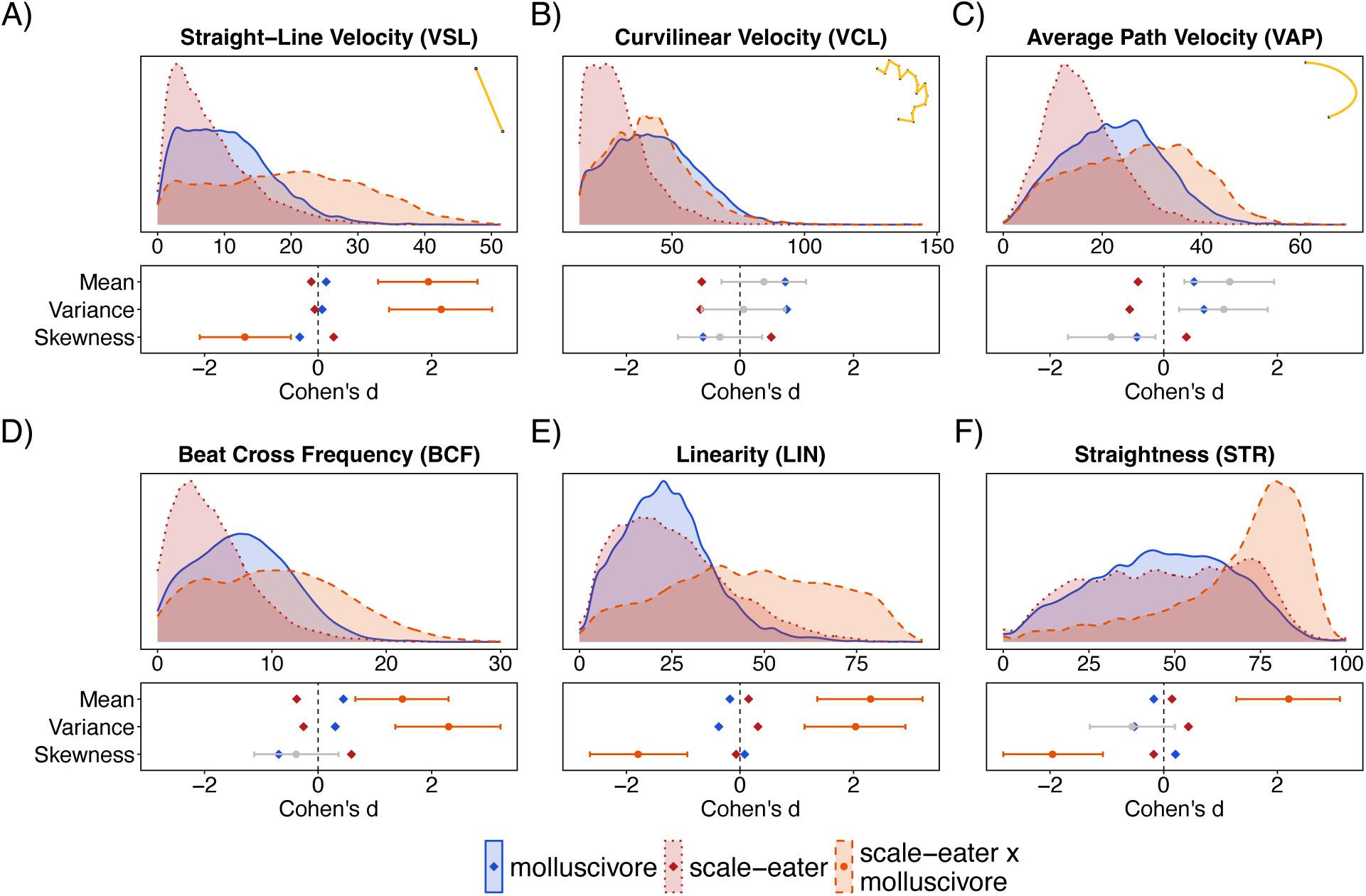
Hybrid sperm kinematics transgress parental distributions. Top subplots show sperm kinematics distributions for hybrids and both parental species after accounting for variation in sample size. Bottom subplots display the directional magnitude of hybrid divergence from the parental complex for mean, variance and skewness. Zero represents the pooled parental species baseline, with each parental *d* marked as well (molluscivore = red, scale-eater = blue). Orange markers represent hybrid *d*, with surrounding stems showing 95% confidence intervals (CIs). Hybrid CIs that don’t overlap with either parental mean are transgressive (marked in orange). Individual sperm distribution analysis confirmed that new hybrid distributions are pervasive and not driven by certain individuals with extreme profiles (Fig. S8). Additionally, complete species analysis shows that generalists and wide-mouths both show similar distributions to parental species and not hybrids (Fig. S9). All kinematic and morphology variable distributions with full moment analyses can be found in Table S3.

Variance in kinematic traits revealed severe phenotypic destabilization within individual hybrid profiles (Fig. 4). The hybrid lineage exhibited transgressive variation in straight-line velocity variance (*d* = 2.167, CI = [1.25, 3.06]), beat-cross frequency variance (*d* = 2.297, CI = [1.36, 3.21]), and linearity variance (*d* = 2.034, CI = [1.13, 2.91]). This manifests as a dramatic flattening and widening of these phenotypic curves in Figure 4.

Skewness confirms that this variation is highly directional, driven by the emergence of extreme phenotypic outliers. We detected transgressive negative shifts in the skew of straightness (*d* = -1.598, CI = [-2.83, -1.07]), linearity (*d* = -1.795, CI = [-2.64, -0.93]), and straight-line velocity (*d* = -1.290, CI = [-2.08, -0.48]) (Fig. 4). Rather than producing a uniform pool of trajectories around optima, hybrid distributions show drastic increases in variation and an unexpected cluster of high straightness, high linearity sperm.

### Trait Correlation Matrices Reveal Unique Increased Hybrid Kinematic Integration

Species-level morphological and kinematic correlation matrices showed large variation in trait correlations across variables (Fig. S10). Integration level (*V_rel_*) did not differ significantly for sperm morphology (*F*_4,48_ = 1.00, *P* = 0.417; Fig. 5A). Kinematic *V_rel_*, however, was significantly different among species (*F*_4,52_ = 4.30, *P* = 0.004; Fig. 5B). Hybrid integration was significantly elevated above that of both parental species (molluscivore: -0.0678 ± 0.0187, *P* = 0.006; scale-eater: 0.0655 ± 0.0180, *P* = 0.006) indicating that hybrid sperm have kinematic variance channeled in fewer multivariate axes than would be expected of intermediate hybrid integration (Fig. 4B; Table S4).

**Fig. 5:**
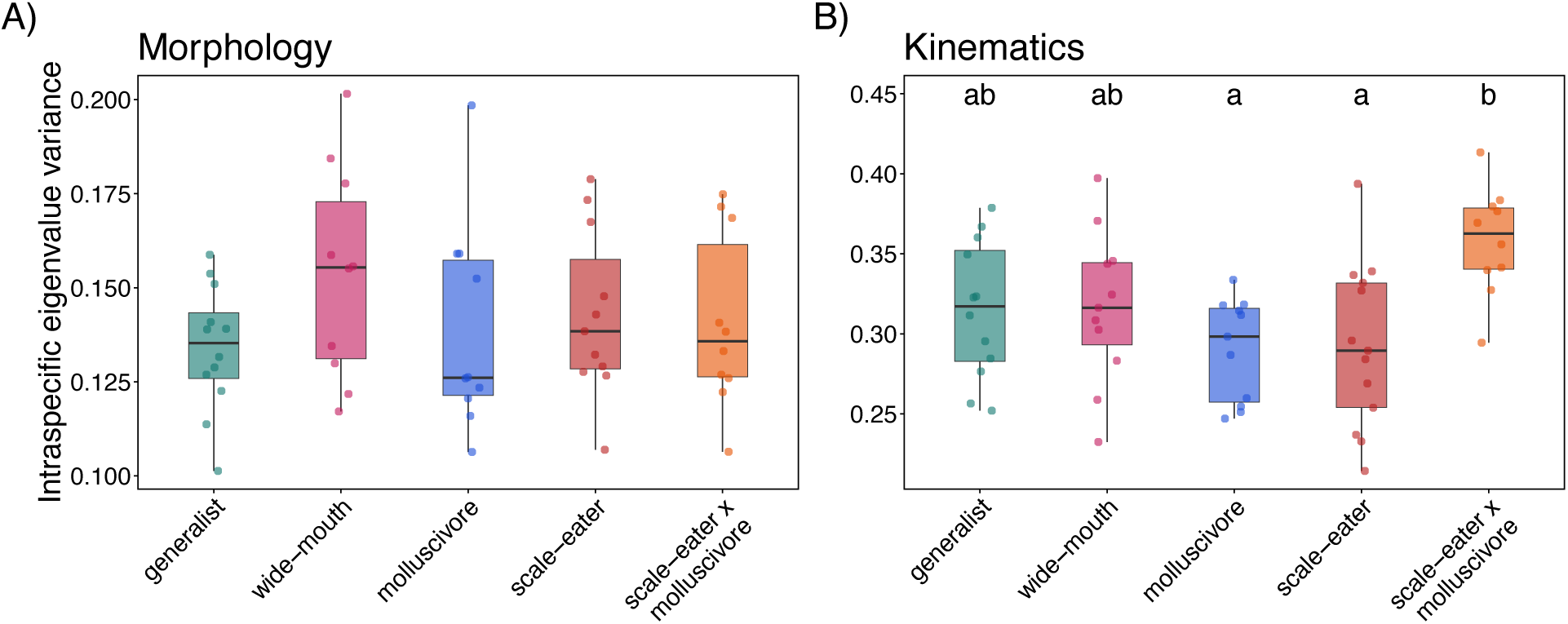
Hybrid sperm show elevated relative eigenvalue variance (*V_rel_*) for kinematic relative to parental species. Boxplots show *V_rel_* calculated from male-level correlation matrices for (A) morphology traits and (B) kinematic traits, with each point representing one individual. Higher eigenvalue variance means trait correlations are concentrated along fewer multivariate axes, reflecting stronger trait integration. Species that do not share a letter grouping significantly differ.

Overall kinematic correlation architecture differed significantly across species (*F*_4,52_ = 5.31, R^2^ = 0.29, *P* < 0.001). Homogeneity of multivariate dispersion across groups (*F*_4,52_ = 0.15, *P* = 0.961) confirmed that these results are not the product of variance differences. Post-hoc pairwise PERMANOVAs revealed that purebred lineages share a highly conserved correlation structure, with no significant differences observed in any purebred–purebred comparison (all *P* > 0.10, Fig. 6A-B). In contrast, hybrid correlation structures differed significantly from every purebred species (all *P* < 0.03; Fig. 6A-B).

**Fig. 6:**
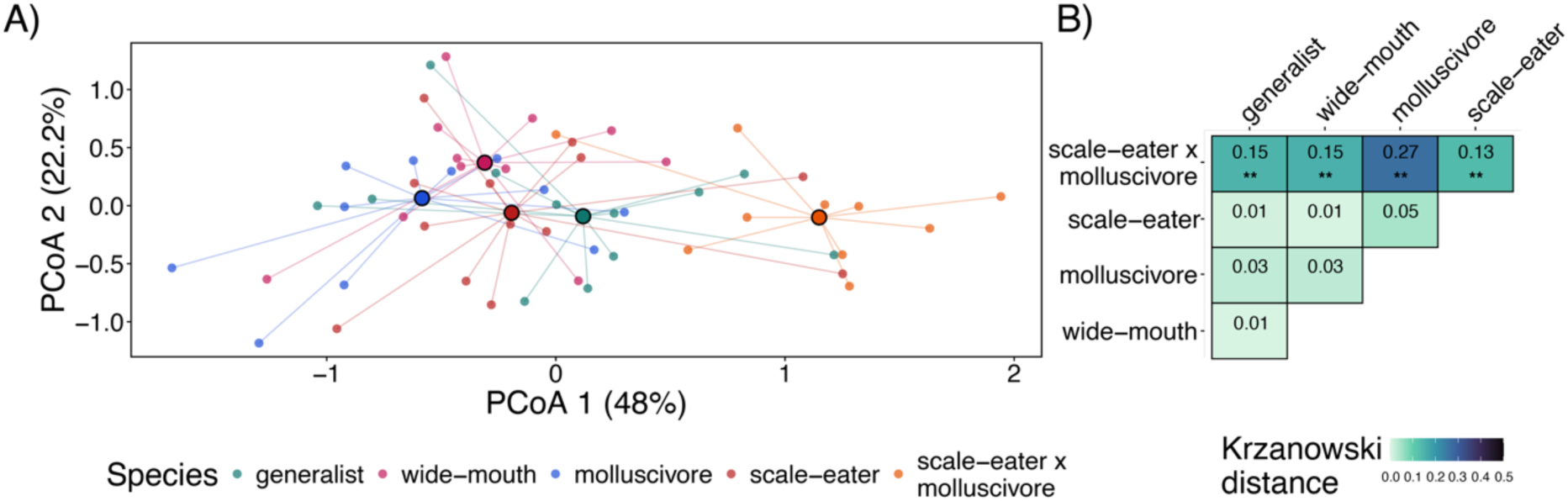
Hybrid kinematic integration is different from all non-hybrid lineages. (A) Principal Coordinate Analysis (PCoA) based on individual Fisher *z*-transformed kinematic correlation matrices. Small points represent male-level kinematic integration matrices connected by line segments to their respective species centroids (large, outlined circles). Percentages on axes denote the proportion of total multivariate variance explained by PCoA 1 and PCoA 2. (B) Heatmap of pairwise Krzanowski distances quantifying subspace divergence across leading principal components (k = traits/2). Lower values (white-teal) represent higher subspace overlap, whereas higher values (blue-black) represent greater matrix divergence. True matrix divergence for this analysis is measured on a scale from 0 to 4, but color range was restricted to better show relative differences. Asterisks indicate significant adjusted differences from post-hoc pairwise PERMANOVAs (* = *P* < 0.05, ** = *P* < 0.01).

Subspace alignment analyses confirmed that hybrid sperm integration operates along structurally distinct axes of variation. Pairwise Krzanowski distances among purebred species were minimal and non-significant (distances = 0.01–0.05; Fig. 6B), indicating nearly identical primary subspace orientation across purebred lineages. Krzanowski distances between hybrids and species were significantly elevated across all comparisons (distances = 0.13–0.27; Fig. 6B). Ordination via PCoA reflected this isolation, with PCoA 1 (48.0% variance) clearly separating the hybrid integration centroid from the closely grouped purebred centroids (Fig. 6A). These results demonstrate that the shift in hybrid integration (*V_rel_*) does not represent a uniform, across-the-board elevation of shared trait correlations, but rather a reorganization of variation within swimming integration architecture.

Morphological correlation architecture also differed significantly across species (*F*_4,52_ = 1.72, *R^2^* = 0.13, *P* = 0.003). However, pairwise comparisons revealed that these differences were driven by morphological differences among non-hybrid species (molluscivore scale-eater divergence: *F* = 2.80, *R^2^* = 0.128, *P* = 0.02) that are consistent with morphological divergence in Figure 2. Krzanowski distances showed that species generally clustered closely with significant subspace divergence only occurring at the outer edges of the species cluster (Fig. S11).

## Discussion

We examined sperm traits across SSI pupfish and their hybrids to assess rapid natural gametic divergence and signs of developing reproductive barriers. We found rapid divergence in sperm morphology and kinematics among sympatric SSI pupfishes. Hybridization between the molluscivore and the scale-eater generated marginal differences in morphology but created transgressive hyper-linear sperm with drastically altered kinematic distributions and kinematic integration patterns.

### Species Divergence in Sperm Morphology and Velocity

Despite the SSI pupfish radiation being ∼10,000 years old (Richards & Martin, 2017), our analysis confirmed substantial divergence in sperm morphology and kinematics among species. Significant differences in midpiece area, flagellum length, and sperm velocity indicate that reproductive traits have diverged alongside the ecological diversification that characterizes this system. Rapid divergence in sperm morphology is common in systems experiencing strong postcopulatory sexual selection, where sperm traits are expected to respond quickly to selection acting on fertilization performance (Birkhead & Pizzari, 2002; Kustra & Alonzo, 2023; Lifjeld, Cramer, et al., 2025; Lüpold et al., 2009; Pitnick et al., 2009; Poignet et al., 2022; Rowe et al., 2015; Tourmente et al., 2011). This selection may reflect divergent life history and demographic constraints within the radiation. Generalists and molluscivores maintain higher relative abundances (Martin & Wainwright, 2013a), potentially driving intense male-male competition where selection primarily optimizes sperm velocity (Fig. 3A). In contrast, scale-eaters occur at much lower population densities (Martin & Wainwright, 2013a) likely reducing sperm competition. Larger midpieces found in scale-eaters and wide-mouths may be a pleiotropic consequence of both species’ independent shifts to specialized scale-eating diets (Fig. 2B, C; Richards & Martin, 2022).

Species also differed substantially in intra-individual sperm trait coefficient of variation (CV). Among non-hybrids, the most prominent pattern was that scale-eaters exhibited higher flagellum length CV relative to both molluscivores and generalists (Fig. 2F). Because intra-individual variation in sperm dimensions directly reflects the strength of stabilizing selection (Immler et al., 2011; Kleven et al., 2008), this elevated CV points to relaxed selection on flagellum length in scale-eaters. While this variation could stem from genetic drift in small founder populations or divergent mating conditions (common in pupfish; Leiser et al., 2015; Leiser & Itzkowitz, 2003, 2004), it seems most consistent with demographic relaxed selection where the extreme rarity of scale-eaters (<2% relative abundance; Martin & Wainwright, 2013a) likely reduces male-male encounters during spawning, weakening the postcopulatory competition that tightly standardizes flagellum length in high-density generalists and molluscivores (Calhim et al., 2007).

### Hybridization Alters Sperm Kinematic Profiles

We found that hybrid sperm exhibit a transgressive hyper-linear phenotype with elevated means in linear kinematic variables relative to non-hybrids (Fig. 4A). This hybrid distribution, influenced by the hyper-linear transgressive phenotype, could be the result of problems in spermatogenesis that arise from hybridization. In teleost fishes, successful fertilization requires sperm to actively navigate osmotic gradients and execute chemotactic reorientations to enter the egg micropyle, a process governed by calcium-dependent switching between symmetric flagellar propagation and asymmetric bending (Alvarez et al., 2014; Cosson et al., 2008; Yanagimachi et al., 2017). This requires sperm to not only swim fast, but also to modulate their curvature and have control over their waveform (Parast et al., 2023). This may mean that the lower levels of integration found in parental species could allow greater kinematic flexibility (Klingenberg, 2014; Pigliucci, 2003). In contrast, hybrid sperm with increased eigenvalue variance may be constrained in modulating their direction, limiting their functional flexibility (Gianoli & Palacio-López, 2009). Consequently, these kinematic transgressions may actually be a byproduct of hybrid breakdown and impair successful sperm performance (McGirr & Martin, 2019; Rieseberg et al., 1999; Stelkens & Seehausen, 2009). This interpretation aligns with known disruptions to spermatogenesis (Hunnicutt et al., 2025; Larson et al., 2017, 2022), suggesting that transgressive hybrid phenotypes could contribute to reproductive isolation between these species.

This kinematic disruption is not a byproduct of morphological divergence, given the lack of transgression in sperm morphology and morphological integration architecture (Figs. 2, S5, S11). Instead, multivariate subspace alignment analyses (Fig. 6) revealed that hybridization produces distinct swimming integration patterns that do not reflect the conserved architectures of the entire radiation. This pattern is reflected in SSI cranial morphology as well, where hybrid integration differs from parental patterns (Chan et al., 2024). These distinct hybrid integration architectures suggest that genetic incompatibilities disrupt the genetic pathways of spermatogenesis (Debat & David, 2001; Rieseberg et al., 1999). To assess whether these incompatibilities directly impact fitness, competitive in vitro fertilization trials could evaluate whether these altered swimming dynamics translate to reduced fertilization success, thereby confirming this transgression as detrimental (Gasparini et al., 2010; Yanagimachi et al., 2017). Ultimately, our findings demonstrate that hybridization between recently diverged species disrupts sperm motility and integration patterns, reinforcing this phenotype as a potential contributor to reproductive isolation.

## Conclusion

While hybrid sperm sterility is a common postzygotic barrier between deeply diverged species, it remains unclear whether fertile hybrids between recently diverged species differ in more subtle ways. We find that, despite similar sperm motility and integration patterns in both parental species, hybridization between recently diverged species (∼10,000 years ago) generates a transgressive, hyper-linear motility phenotype and substantial reorganization of kinematic covariance structures and trait distributions. Because morphology was unaffected, these changes seem to be caused by a breakdown in the genetic, regulatory, or physiological control of motility. Ultimately, these transgressive kinematic signatures demonstrate how hybridization can reorganize fine-tuned gametes, providing crucial insight into the formation of intrinsic postzygotic barriers during the early stages of speciation.

## Author Contributions

OG and MCK conceived and designed the study with input from CHM. OG conducted the experiment. OG and MCK analyzed the data. OG drafted the initial manuscript, CHM and MCK critically revised the manuscript. All authors approve of the publication of this article.

## Acknowledgements

We thank the Gerace Research Center and Troy Day for logistical support in the Bahamas, and the government of the Bahamas BEST commission for permission to collect and export samples (Permit No. PPF/DGOPA-001/20). We thank Suzanne Alonzo for the microscope and software used for sperm motility analysis.

## Supplemental Information for

### Supplemental Information Includes

Supplemental Methods: Morphology-Kinematic relationships

Supplemental Results: Morphology-Kinematic relationships

Figures S1-S11

Tables S1-S5

### Supplemental Methods: Morphology-Kinematic relationships

We compared species differences in morphology-kinematic relationships by averaging sperm metrics at the per-male level and merging morphology metrics with the corresponding males mean velocity value. We conducted separate linear models for each morphology-velocity combination (head to tail ratio was included as a morphology trait in this analysis because of its relevance to sperm velocities; Humphries et al., 2008), using type III ANOVAs from the car package to assess significance (Fox et al., 2026). We tested these interaction models across the 4 species and the hybrid group.

### Supplemental Results: Morphology-Kinematic relationships

All species showed similar, weak morphology-kinematic relationships (Fig. S6). Flagellum length was a positive predictor of sperm velocity across all species (*F*_1,48_ = 15.77, *P* < 0.001, *ƞp²* = 0.25) while head area (*F*_1,48_ = 11.90, *P* = 0.001, *ƞp²* = 0.20) and head-tail ratio (*F*_1,48_ = 22.80, *P* < 0.001, *ƞp²* = 0.32) were both strong negative predictors of velocity. The relationship between sperm morphology and velocity did not differ among species or in hybrids (Fig. S6).

## Supplemental Information

**Fig. S1:**
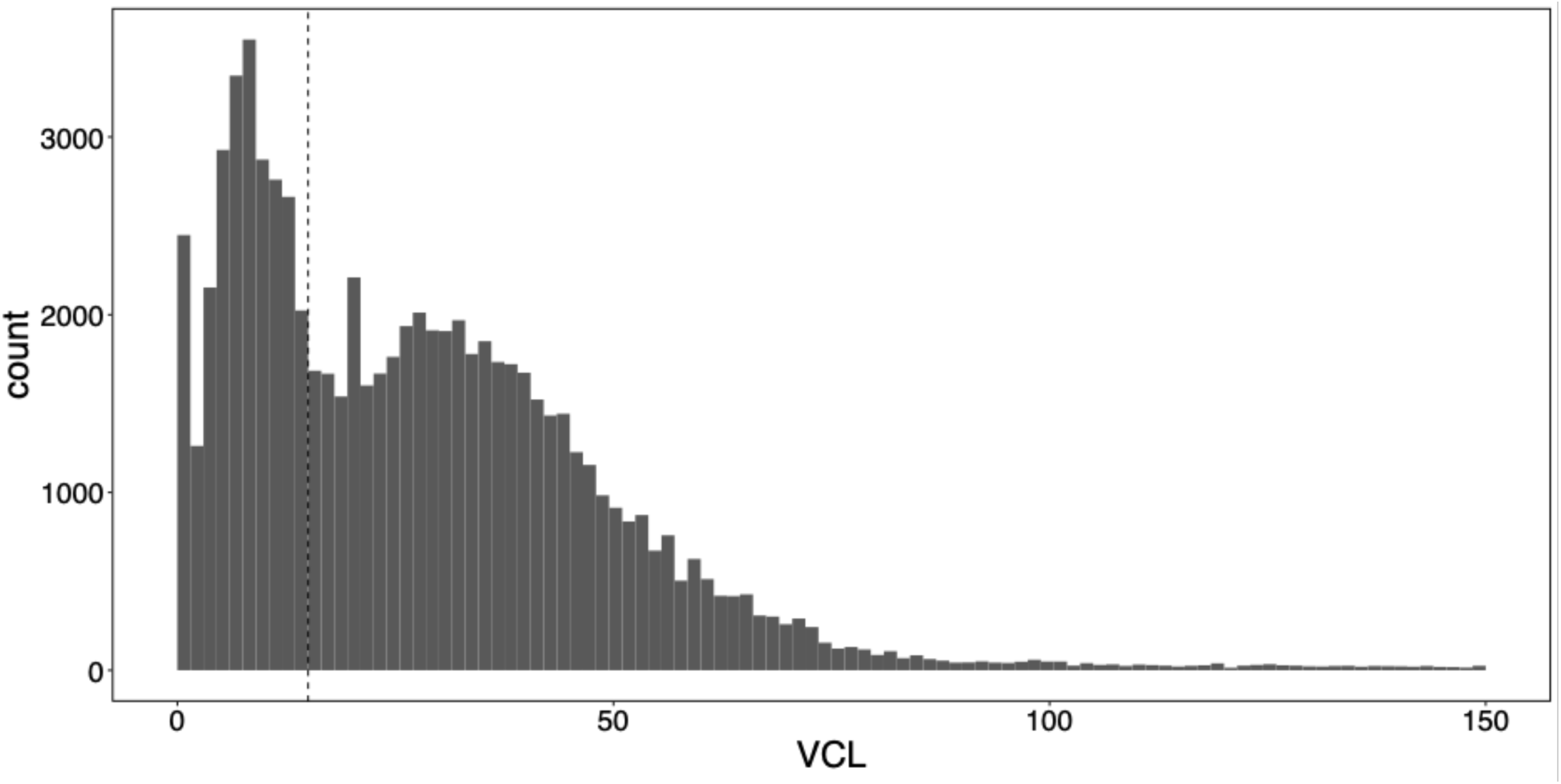
Distribution of curvilinear velocity (VCL) of all sperm measured demonstrating bimodal distribution. Distribution was used to filter non-motile sperm from data set as minimal total movement incurs larger error with rate statistics such as STR, LIN, and WOB. Additionally, minimal disruptions to recording can cause non-motile sperm to appear within 0-10 VCL that cause the second peak of this bimodal distribution. Dashed line indicates the 15 VCL cutoff that was used when filtering.

**Fig. S2:**
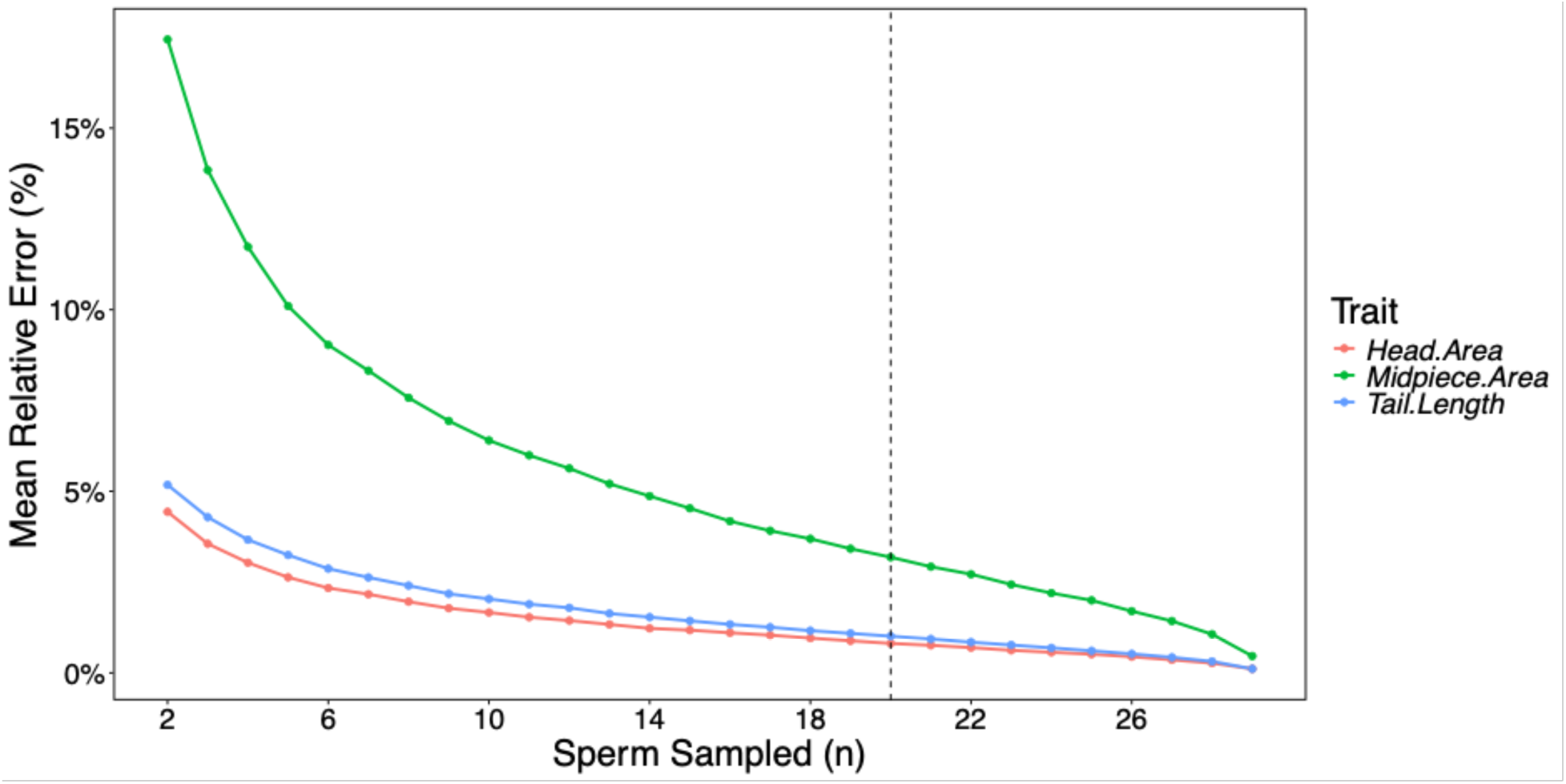
Mean relative estimation error exhibits diminishing returns above n = 20 sperm measurements per sample, placing this threshold well past the curve’s primary inflection point. The midpiece displays higher relative error across all sample sizes because its small absolute dimensions magnify equivalent manual measurement imprecision compared to head area and flagellum length. At n = 20, mean relative error falls below 5% for all morphometric traits, confirming that measuring 20 sperm per sample provides sufficient precision.

**Fig. S3:**
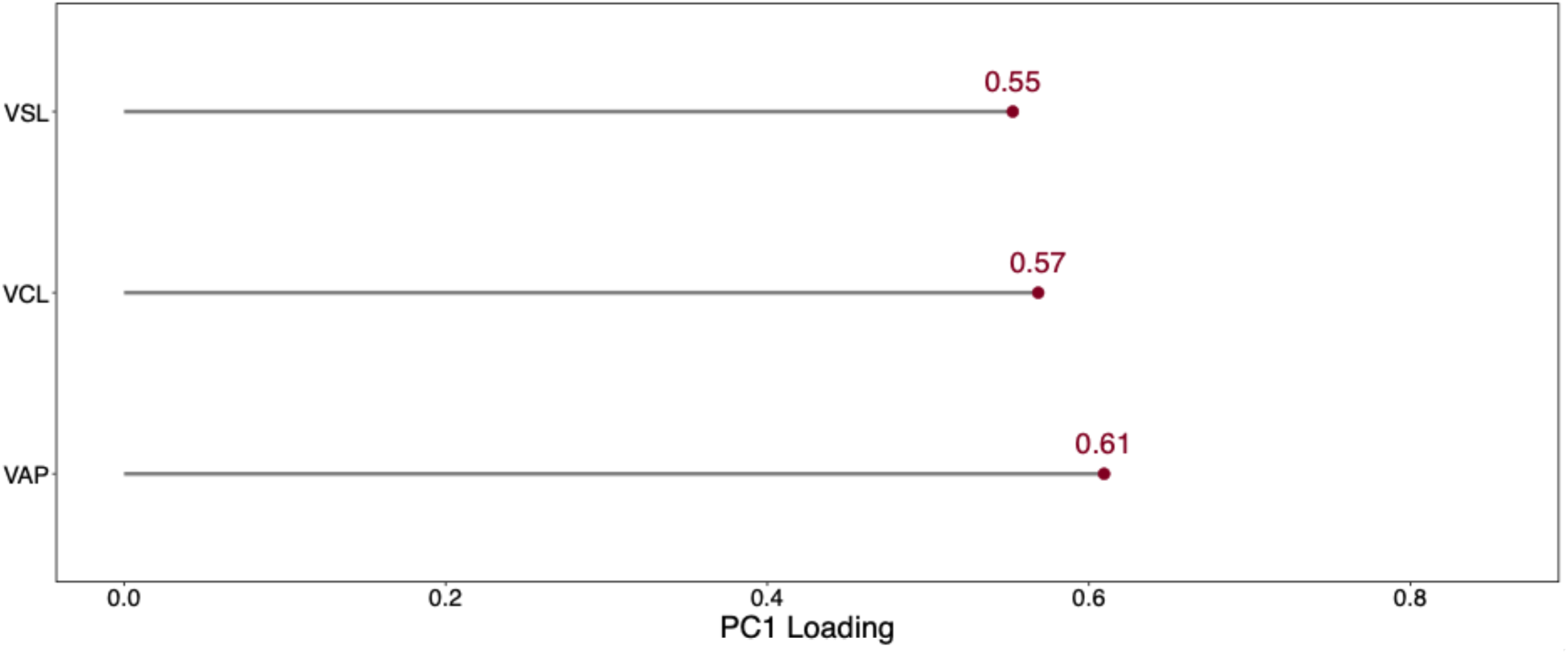
Principal component loadings defining the composite sperm velocity axis. The plot displays the variable loadings (eigenvectors) on the first principal component (PC1) derived from a Principal Component Analysis (PCA) of straight-line velocity (VSL), curvilinear velocity (VCL), and average path velocity (VAP). Because PC1 successfully captures shared variance amongst all three traits, individual PC1 scores were extracted and utilized as a single, composite velocity component.

**Fig. S4:**
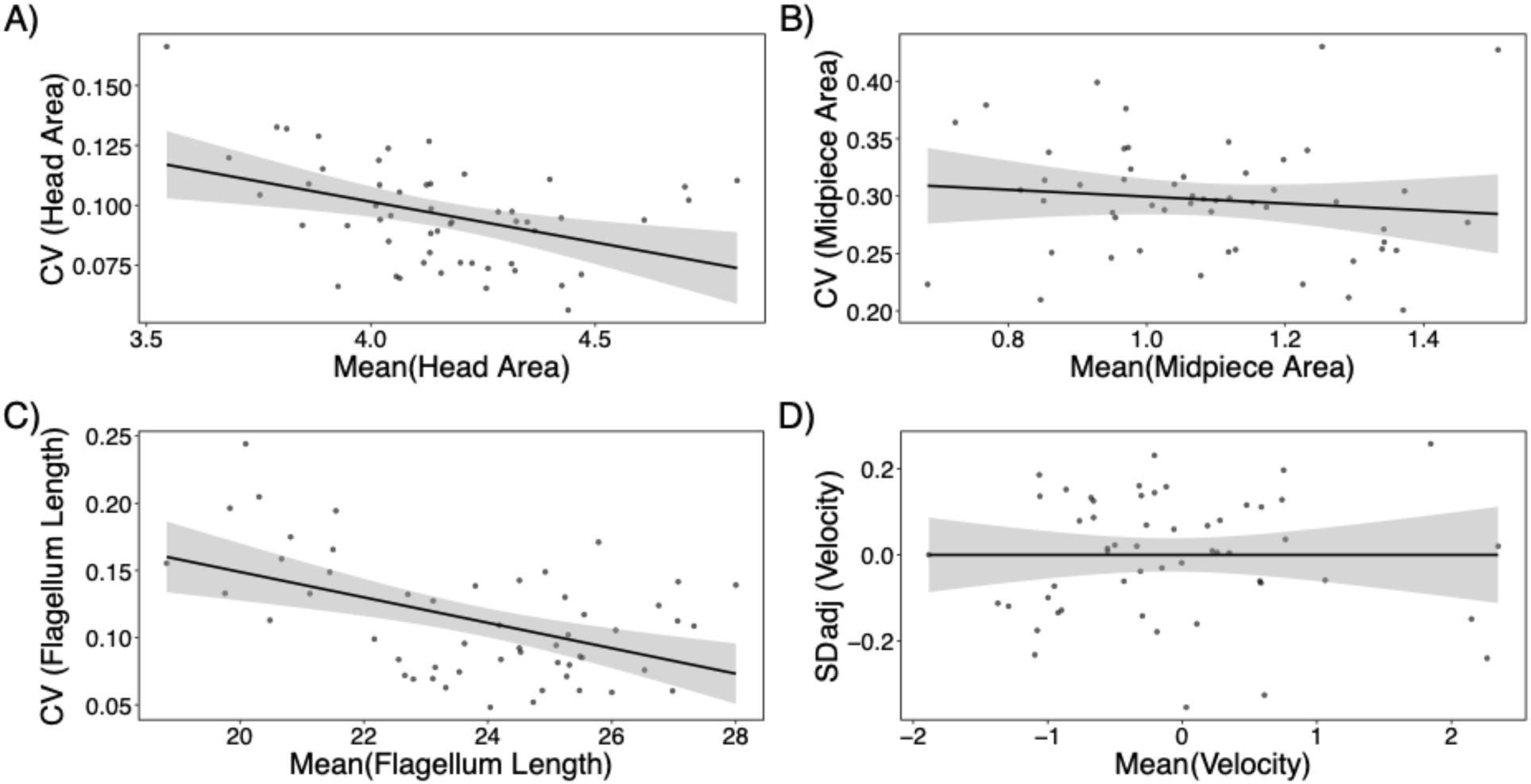
Relationship between mean and mean-adjusted variance metrics shows removal of mean-variance scaling. Scatter plots displaying the intra-individual mean versus the intra-individual variance metric used in Fig. 2 and Fig. 3 for (A) head area, (B) midpiece area, (C) flagellum length, and (D) swimming velocity. Each point represents an individual male. The flat linear regression lines visually demonstrate that the CV and SD_adj_ metrics successfully decouple trait variance from absolute trait size, eliminating mean-scale dependency across all measured traits prior to statistical analysis.

**Fig. S5:**
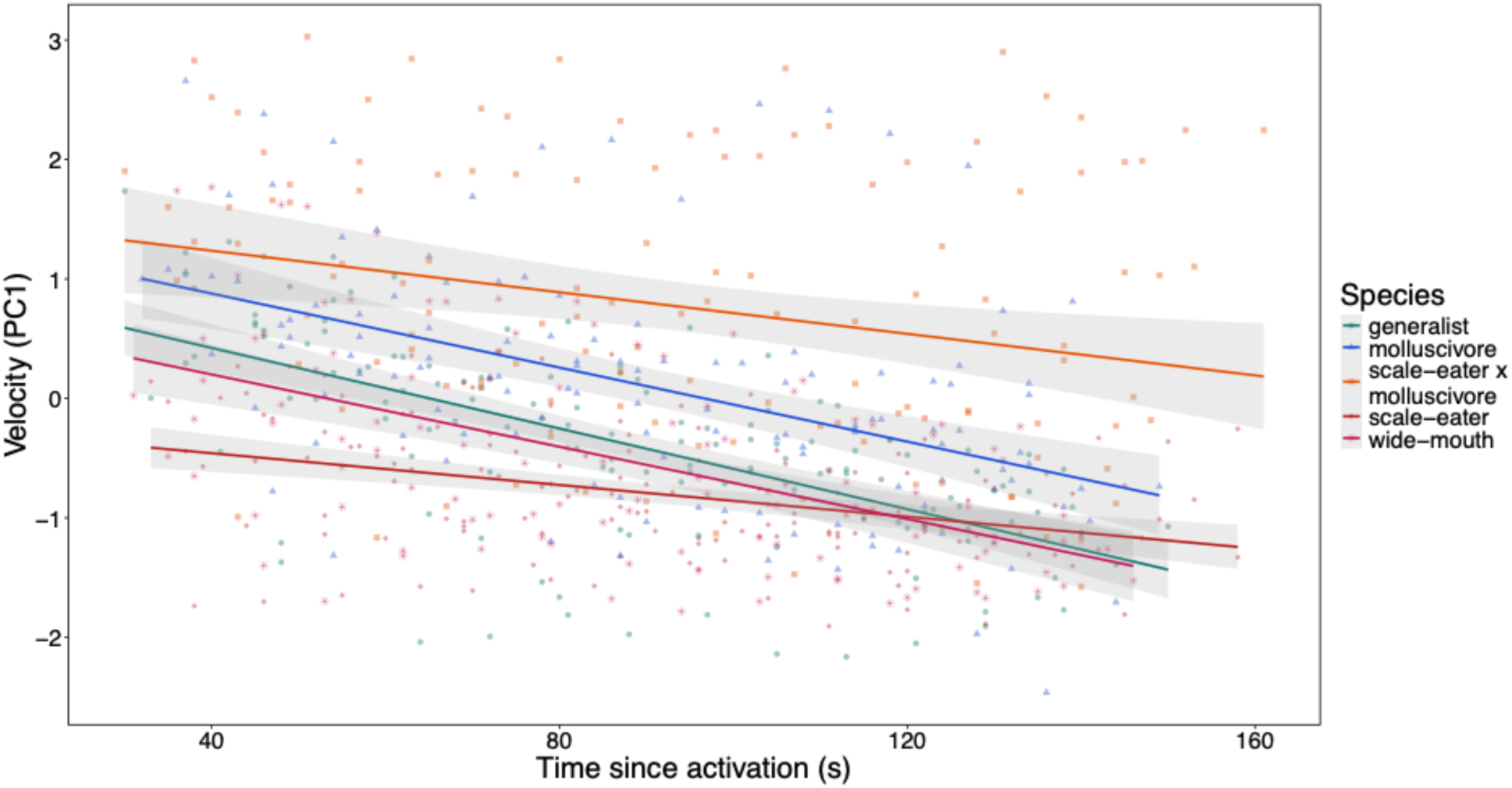
Hybrids do not show abnormal reduction in longevity despite increased velocity metrics. Plot compares combined velocity metric (Fig. S3) and time since activation with each species as a separate group. The hybrid lineage sustains elevated velocities throughout the sampled timeframe.

**Fig. S6:**
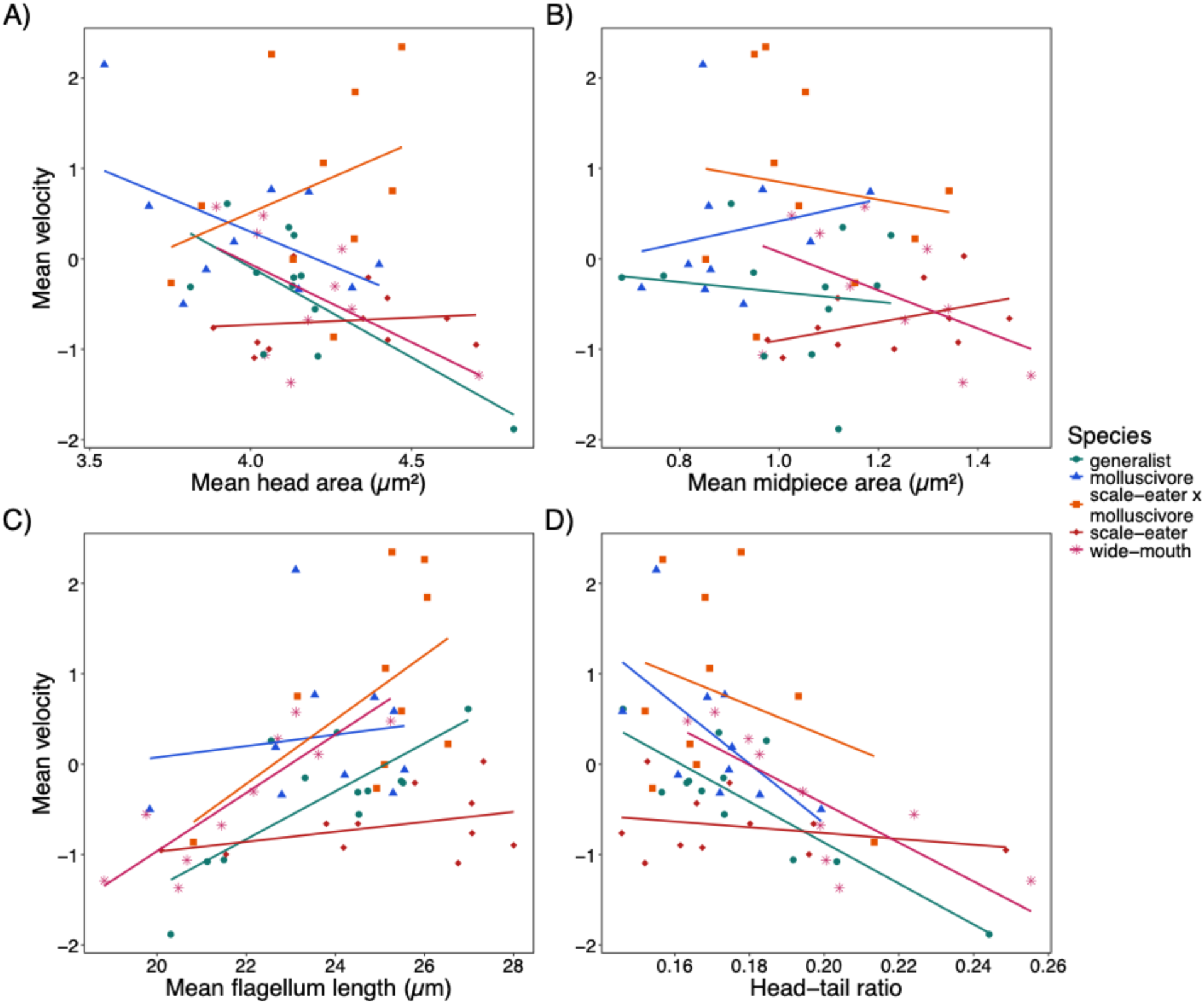
Species show similar morphology-velocity relationships. Relationships between mean sperm velocity and (A) mean head area, (B) mean midpiece area, and (C) mean flagellum length across species. Points represent male-level means and lines indicate species-specific linear regressions for each morphological trait. Mean velocity value was calculated as the dominant component axis of a PCA on the individual sperm variables VSL, VCL, and VAP.

**Fig. S7:**
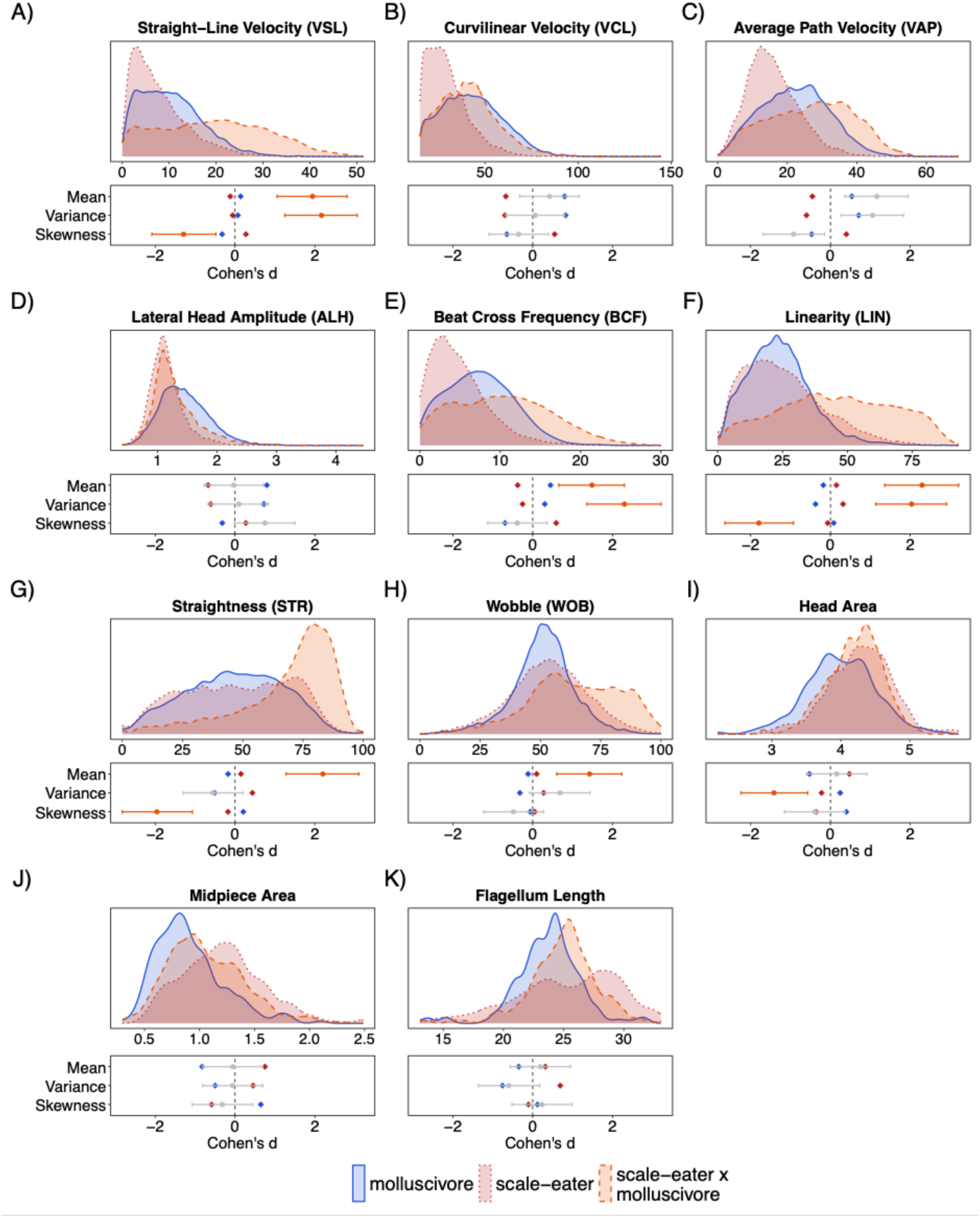
Complete morphology and kinematic distributions amongst hybrids and parental species. Top subplots show distributions of individual sperm across morphology (A-C) and kinematic (D-K) traits after accounting for variation among individuals. Bottom subplots display the directional magnitude of hybrid (C. brontotheroides × C. desquamator) divergence from the parental complex for mean, variance, skewness, and kurtosis. Zero represents the pooled parental species baseline, calculated from the collective mean of both parental species while Cohen’s d-statistic represents the directional magnitude of difference in the hybrid, expressed in standard deviation units. Markers and connecting stems colored in pink highlight transgressive traits where the hybrids significantly differed from both parental species. Distributions and variance measures capture a marked widening and flattening of many hybrid distributions (e.g. VSL, VAP, LIN, and BCF) as well as unusual shift in skew for STR and LIN and leptokurtic clustering at the upper phenotypic boundary in STR.

**Fig. S8.**
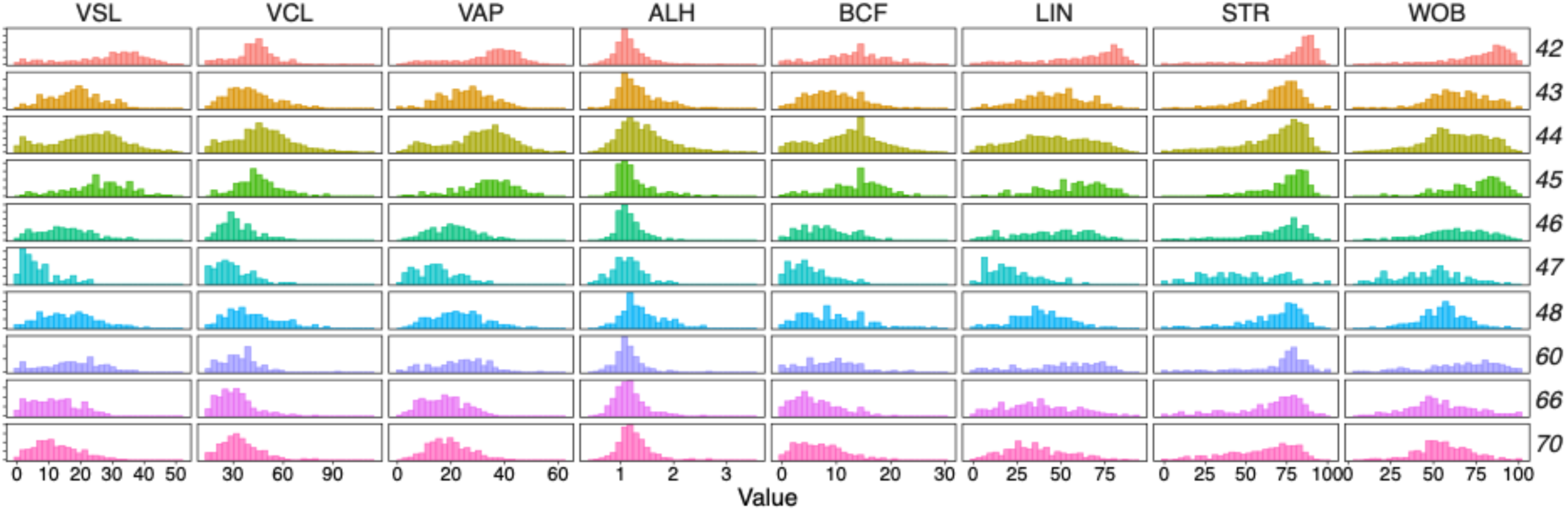
Distribution of all hybrid individual kinematics showing pervasive distribution change. Numbers on the right side indicate ID number for hybrid individuals and distributions are histograms of sperm within that individual. Distributions all generally align showing consistent disruption of purebred species distributions rather than single individual influence

**Fig. S9:**
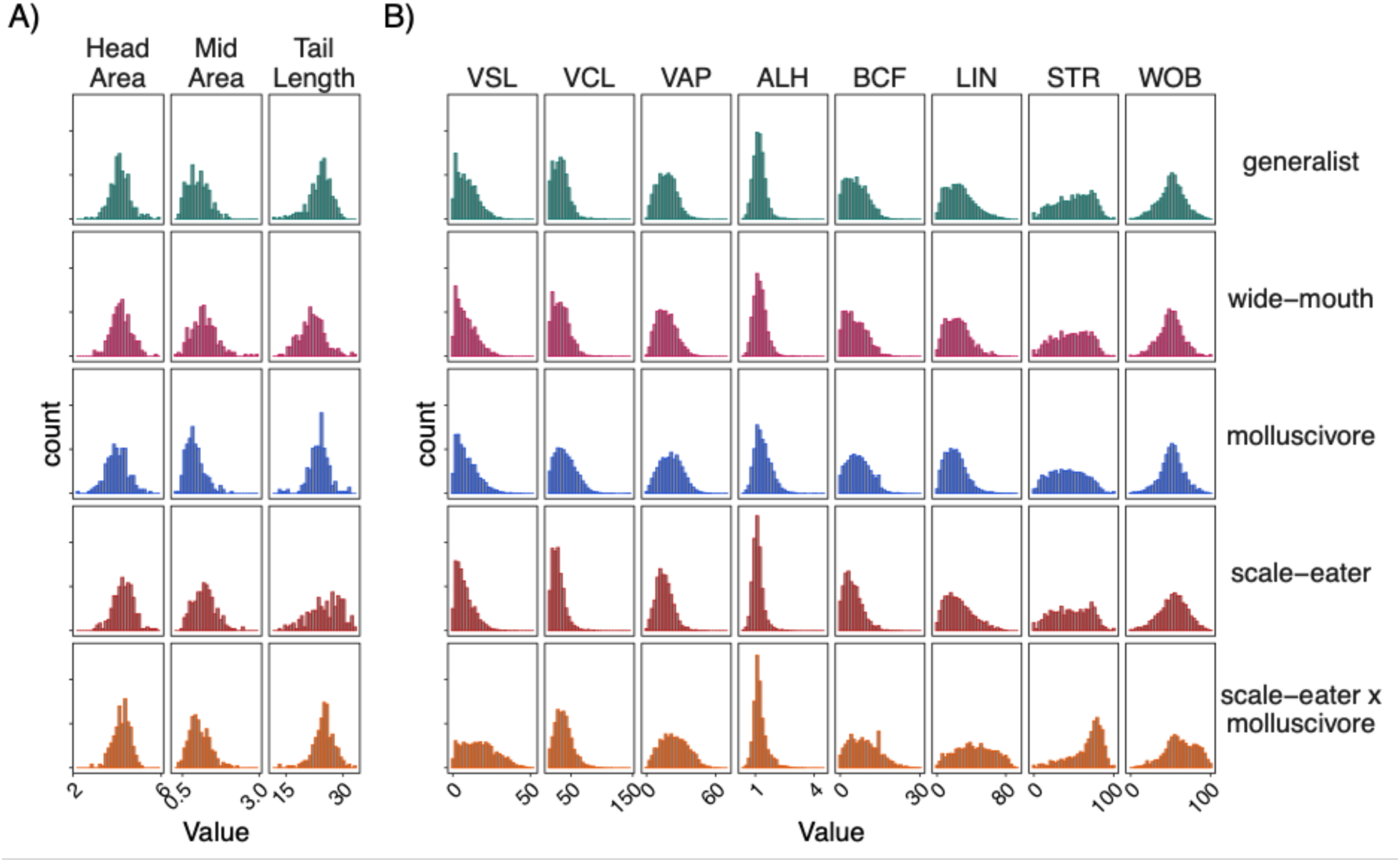
Trait distributions for all purebred species mirror those of parental species. Shows detailed distributions for (A) morphology and (B) kinematics for all species. Generalists (*C.* variegatus) and wide-mouths (*C. sp.* ‘wide-mouth’) both show similar distributions to parentals shown in Fig. 4. Histograms additionally show further detail in exact distributions that smoothed curves in Fig. 4 and Fig. S7 suppress.

**Fig. S10:**
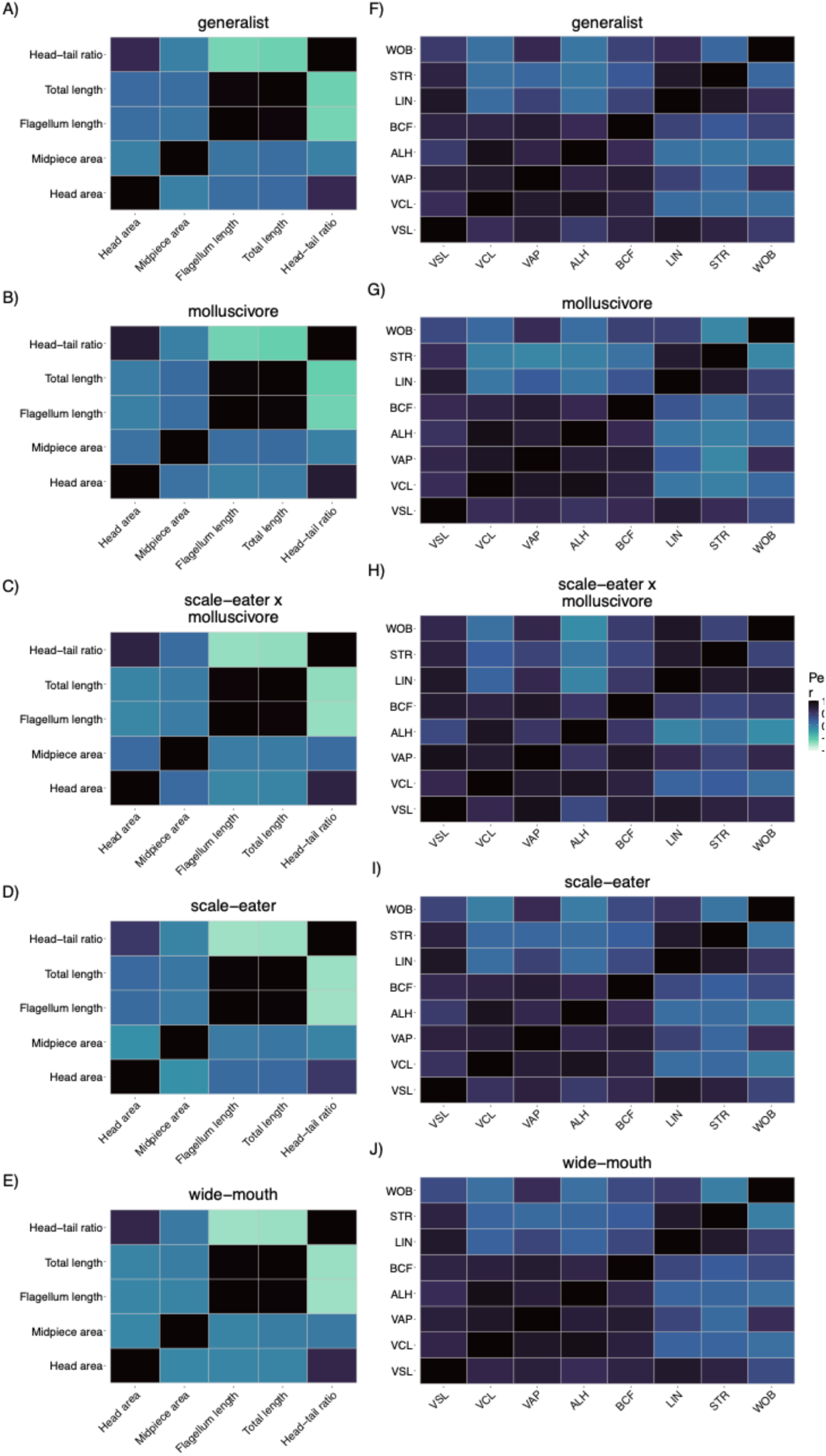
Lineage-specific pairwise correlation matrices for sperm morphological and kinematic traits. Green areas show positive correlation while orange areas show negative correlation. Correlation structures across purebred species remain highly conserved in both trait modules, however hybrid sperm have less white in their kinematic heatmap, supporting higher levels of kinematic integration. See Figure 1B for description of kinematic variables.

**Fig. S11:**
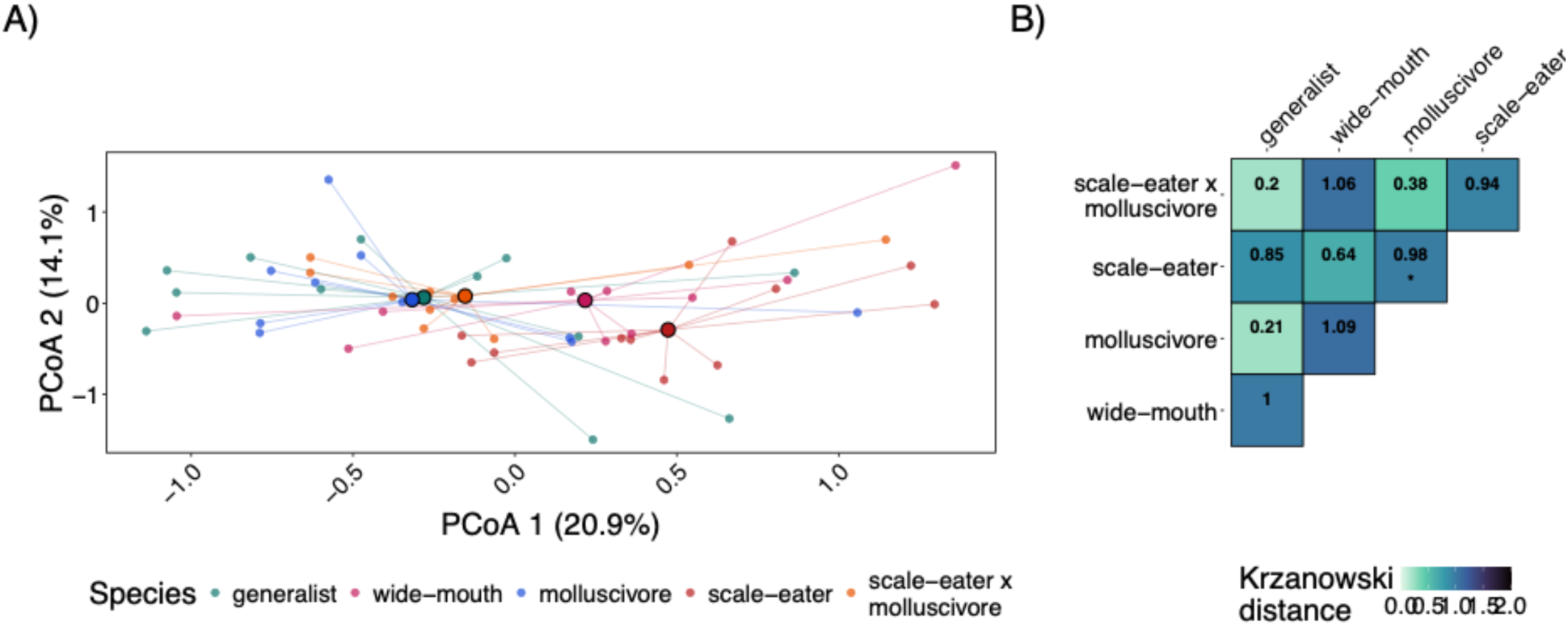
Morphological integration shows intermediate hybrid and complete species clustering. (A) Principal Coordinate Analysis (PCoA) based on individual Fisher *z*-transformed morphological correlation matrices. Small points represent male-level morphological integration matrices connected by line segments to their respective species centroids (large, outlined circles). Percentages on axes denote the proportion of total multivariate variance explained by PCoA 1 and PCoA 2. (B) Heatmap of pairwise Krzanowski distances quantifying subspace divergence across leading principal components (k = traits/2). Lower values (darker green) represent higher subspace overlap, whereas higher values represent greater matrix orientation divergence. Asterisks indicate adjusted significance from post-hoc pairwise PERMANOVAs (* = *P* < 0.05, ** = *P* < 0.01).

**Table S1:** Pairwise species comparisons for sperm morphological trait means and variances. Estimated contrast differences (Difference), standard errors (SE), Satterthwaite degrees of freedom (df), *t*-statistics, and adjusted *P*-values for pairwise comparisons of flagellum length, head area, and midpiece area between species. Trait means were evaluated using linear mixed-effects models (LMMs) with individual ID specified as a random intercept to account for repeated measures per male. Trait variances were evaluated using linear models (LMs). Pairwise contrasts (Species A vs. Species B) were calculated using estimated marginal means (emmeans) with Tukey-adjusted *P*-values for multiple comparisons. Bold rows indicate significant

| Trait | Metric | Species A | Species B | Difference | SE | df | t | P |
| --- | --- | --- | --- | --- | --- | --- | --- | --- |
| Flagellum | Mean | generalist | molluscivore | 0.02 | 0.86 | 48.1 | 0.02 | 1.000 |
| Flagellum | Mean | generalist | scale-eater x molluscivore | -1.12 | 0.86 | 47.7 | -1.30 | 0.690 |
| Flagellum | Mean | generalist | scale-eater | -1.39 | 0.84 | 48.3 | -1.65 | 0.471 |
| Flagellum | Mean | generalist | wide-mouth | 1.92 | 0.86 | 48.0 | 2.23 | 0.187 |
| Flagellum | Mean | molluscivore | scale-eater x molluscivore | -1.14 | 0.90 | 47.8 | -1.27 | 0.713 |
| Flagellum | Mean | molluscivore | scale-eater | -1.41 | 0.88 | 48.3 | -1.60 | 0.506 |
| Flagellum | Mean | molluscivore | wide-mouth | 1.90 | 0.90 | 48.0 | 2.11 | 0.231 |
| Flagellum | Mean | scale-eater x molluscivore | scale-eater | -0.27 | 0.88 | 47.9 | -0.31 | 0.998 |
| **Flagellum** | **Mean** | **scale-eater x molluscivore** | **wide-mouth** | **3.04** | **0.90** | **47.6** | **3.39** | **0.012** |
| **Flagellum** | **Mean** | **scale-eater** | **wide-mouth** | **3.31** | **0.88** | **48.2** | **3.76** | **0.004** |
| Head | Mean | generalist | molluscivore | 0.15 | 0.11 | 48.1 | 1.38 | 0.644 |
| Head | Mean | generalist | scale-eater x molluscivore | -0.05 | 0.11 | 47.5 | -0.43 | 0.993 |
| Head | Mean | generalist | scale-eater | -0.13 | 0.10 | 48.4 | -1.27 | 0.711 |
| Head | Mean | generalist | wide-mouth | -0.05 | 0.11 | 47.9 | -0.43 | 0.993 |
| Head | Mean | molluscivore | scale-eater x molluscivore | -0.19 | 0.11 | 47.7 | -1.73 | 0.424 |
| Head | Mean | molluscivore | scale-eater | -0.28 | 0.11 | 48.4 | -2.56 | 0.094 |
| Head | Mean | molluscivore | wide-mouth | -0.19 | 0.11 | 48.0 | -1.73 | 0.426 |
| Head | Mean | scale-eater x molluscivore | scale-eater | -0.09 | 0.11 | 47.8 | -0.80 | 0.930 |
| Head | Mean | scale-eater x molluscivore | wide-mouth | 0.00 | 0.11 | 47.5 | 0.00 | 1.000 |
| Head | Mean | scale-eater | wide-mouth | 0.09 | 0.11 | 48.2 | 0.79 | 0.931 |
| Midpiece | Mean | generalist | molluscivore | 0.10 | 0.07 | 48.2 | 1.52 | 0.554 |
| Midpiece | Mean | generalist | scale-eater x molluscivore | -0.04 | 0.07 | 47.2 | -0.66 | 0.965 |
| **Midpiece** | **Mean** | **generalist** | **scale-eater** | **-0.20** | **0.07** | **48.5** | **-3.00** | **0.033** |
| **Midpiece** | **Mean** | **generalist** | **wide-mouth** | **-0.20** | **0.07** | **47.9** | **-2.93** | **0.039** |
| Midpiece | Mean | molluscivore | scale-eater x molluscivore | -0.15 | 0.07 | 47.5 | -2.09 | 0.241 |
| **Midpiece** | **Mean** | **molluscivore** | **scale-eater** | **-0.30** | **0.07** | **48.7** | **-4.35** | **<0.001** |
| **Midpiece** | **Mean** | **molluscivore** | **wide-mouth** | **-0.30** | **0.07** | **48.1** | **-4.26** | **<0.001** |
| Midpiece | Mean | scale-eater x molluscivore | scale-eater | -0.16 | 0.07 | 47.7 | -2.24 | 0.183 |
| Midpiece | Mean | scale-eater x molluscivore | wide-mouth | -0.16 | 0.07 | 47.2 | -2.19 | 0.201 |
| Midpiece | Mean | scale-eater | wide-mouth | 0.00 | 0.07 | 48.4 | 0.00 | 1.000 |
| Flagellum | CV | generalist | molluscivore | 0.00 | 0.02 | 48.0 | 0.14 | 1.000 |
| Flagellum | CV | generalist | scale-eater x molluscivore | 0.00 | 0.02 | 48.0 | -0.16 | 1.000 |
| **Flagellum** | **CV** | **generalist** | **scale-eater** | **-0.05** | **0.02** | **48.0** | **-3.23** | **0.018** |
| Flagellum | CV | generalist | wide-mouth | -0.03 | 0.02 | 48.0 | -2.05 | 0.259 |
| Flagellum | CV | molluscivore | scale-eater x molluscivore | -0.01 | 0.02 | 48.0 | -0.29 | 0.998 |
| **Flagellum** | **CV** | **molluscivore** | **scale-eater** | **-0.06** | **0.02** | **48.0** | **-3.22** | **0.019** |
| Flagellum | CV | molluscivore | wide-mouth | -0.04 | 0.02 | 48.0 | -2.09 | 0.239 |
| **Flagellum** | **CV** | **scale-eater x molluscivore** | **scale-eater** | **-0.05** | **0.02** | **48.0** | **-2.93** | **0.040** |
| Flagellum | CV | scale-eater x molluscivore | wide-mouth | -0.03 | 0.02 | 48.0 | -1.81 | 0.382 |
| Flagellum | CV | scale-eater | wide-mouth | 0.02 | 0.02 | 48.0 | 1.08 | 0.817 |
| Head | CV | generalist | molluscivore | -0.01 | 0.01 | 48.0 | -1.77 | 0.403 |
| Head | CV | generalist | scale-eater x molluscivore | 0.02 | 0.01 | 48.0 | 2.28 | 0.170 |
| Head | CV | generalist | scale-eater | 0.00 | 0.01 | 48.0 | 0.21 | 1.000 |
| Head | CV | generalist | wide-mouth | 0.00 | 0.01 | 48.0 | -0.29 | 0.998 |
| **Head** | **CV** | **molluscivore** | **scale-eater x molluscivore** | **0.03** | **0.01** | **48.0** | **3.88** | **0.003** |
| Head | CV | molluscivore | scale-eater | 0.02 | 0.01 | 48.0 | 1.93 | 0.314 |
| Head | CV | molluscivore | wide-mouth | 0.01 | 0.01 | 48.0 | 1.42 | 0.619 |
| Head | CV | scale-eater x molluscivore | scale-eater | -0.02 | 0.01 | 48.0 | -2.03 | 0.266 |
| Head | CV | scale-eater x molluscivore | wide-mouth | -0.02 | 0.01 | 48.0 | -2.46 | 0.118 |
| Head | CV | scale-eater | wide-mouth | 0.00 | 0.01 | 48.0 | -0.48 | 0.989 |
| Midpiece | CV | generalist | molluscivore | -0.02 | 0.02 | 48.0 | -0.75 | 0.944 |
| Midpiece | CV | generalist | scale-eater x molluscivore | 0.00 | 0.02 | 48.0 | -0.12 | 1.000 |
| Midpiece | CV | generalist | scale-eater | 0.01 | 0.02 | 48.0 | 0.58 | 0.977 |
| Midpiece | CV | generalist | wide-mouth | -0.01 | 0.02 | 48.0 | -0.58 | 0.978 |
| Midpiece | CV | molluscivore | scale-eater x molluscivore | 0.01 | 0.02 | 48.0 | 0.60 | 0.974 |
| Midpiece | CV | molluscivore | scale-eater | 0.03 | 0.02 | 48.0 | 1.29 | 0.700 |
| Midpiece | CV | molluscivore | wide-mouth | 0.00 | 0.02 | 48.0 | 0.16 | 1.000 |
| Midpiece | CV | scale-eater x molluscivore | scale-eater | 0.02 | 0.02 | 48.0 | 0.67 | 0.961 |
| Midpiece | CV | scale-eater x molluscivore | wide-mouth | -0.01 | 0.02 | 48.0 | -0.44 | 0.992 |
| Midpiece | CV | scale-eater | wide-mouth | -0.03 | 0.02 | 48.0 | -1.12 | 0.794 |

**Table S2:** Pairwise species comparisons for sperm composite velocity mean and coefficient of variation (CV). Estimated contrast differences (Difference), standard errors (SE), Satterthwaite degrees of freedom (df), *t-*statistics (t), and adjusted *P*-values for pairwise comparisons of composite velocity metric (Fig. 3A) between species. Means were evaluated using linear mixed-effects models (LMMs) with individual ID specified as a random intercept to account for repeated measures per male. Trait variances were evaluated using linear models (LMs). Pairwise contrasts (Species A vs. Species B) were calculated using estimated marginal means (emmeans) with Tukey-adjusted *P*-values for multiple comparisons. Bold rows indicate significant differences between groups (*P* < 0.05).

| Trait | Metric | Species A | Species B | Difference | SE | df | t | P |
| --- | --- | --- | --- | --- | --- | --- | --- | --- |
| Velocity | Mean | generalist | molluscivore | -0.54 | 0.32 | 51.8 | -1.70 | 0.445 |
| **Velocity** | **Mean** | **generalist** | **scale-eater x molluscivore** | **-1.17** | **0.33** | **52.0** | **-3.57** | **0.007** |
| Velocity | Mean | generalist | scale-eater | 0.38 | 0.31 | 52.1 | 1.24 | 0.725 |
| Velocity | Mean | generalist | wide-mouth | 0.02 | 0.32 | 52.2 | 0.07 | 1.000 |
| Velocity | Mean | molluscivore | scale-eater x molluscivore | -0.63 | 0.33 | 51.7 | -1.88 | 0.343 |
| **Velocity** | **Mean** | **molluscivore** | **scale-eater** | **0.92** | **0.31** | **51.8** | **2.95** | **0.037** |
| Velocity | Mean | molluscivore | wide-mouth | 0.56 | 0.33 | 51.9 | 1.73 | 0.426 |
| **Velocity** | **Mean** | **scale-eater x molluscivore** | **scale-eater** | **1.55** | **0.32** | **52.0** | **4.81** | **<0.001** |
| **Velocity** | **Mean** | **scale-eater x molluscivore** | **wide-mouth** | **1.19** | **0.33** | **52.1** | **3.56** | **0.007** |
| Velocity | Mean | scale-eater | wide-mouth | -0.36 | 0.31 | 52.2 | -1.15 | 0.781 |
| Velocity | SDadj | generalist | molluscivore | -0.09 | 0.06 | 48.0 | -1.58 | 0.518 |
| Velocity | SDadj | generalist | scale-eater x molluscivore | -0.10 | 0.06 | 48.0 | -1.68 | 0.454 |
| Velocity | SDadj | generalist | scale-eater | 0.00 | 0.06 | 48.0 | 0.04 | 1.000 |
| Velocity | SDadj | generalist | wide-mouth | -0.07 | 0.06 | 48.0 | -1.19 | 0.759 |
| Velocity | SDadj | molluscivore | scale-eater x molluscivore | -0.01 | 0.06 | 48.0 | -0.10 | 1.000 |
| Velocity | SDadj | molluscivore | scale-eater | 0.09 | 0.06 | 48.0 | 1.58 | 0.514 |
| Velocity | SDadj | molluscivore | wide-mouth | 0.02 | 0.06 | 48.0 | 0.38 | 0.996 |
| Velocity | SDadj | scale-eater x molluscivore | scale-eater | 0.10 | 0.06 | 48.0 | 1.69 | 0.452 |
| Velocity | SDadj | scale-eater x molluscivore | wide-mouth | 0.03 | 0.06 | 48.0 | 0.48 | 0.989 |
| Velocity | SDadj | scale-eater | wide-mouth | -0.07 | 0.06 | 48.0 | -1.20 | 0.752 |

**Table S3:** Standardized effect sizes and transgressive segregation analysis across sperm kinematic and morphological moments. Standardized phenotypic differences (Cohen’s *d*) and 95% confidence intervals (CIs) calculated for scale-eater x molluscivore hybrids relative to the pooled parental baseline across three distributional moments (Mean, Variance, and Skewness) for all measured kinematic and morphological traits. To evaluate transgression on a unified scale without relying on null hypothesis significance testing or pairwise *P*-value thresholding, parental species means (d_M: molluscivore; d_P: scale-eater) were mapped onto this same coordinate system as standard deviation offsets from the parental baseline average. A moment was formally classified as transgressive (Transgressive_Flag = TRUE and bolded in table) if and only if both parental values (d_M and d_P) fell outside the hybrid 95% confidence interval. This criterion ensures that transgressive designations reflect genuine phenotypic expansion beyond either parental boundary.

| trait | moment | Cohens_d | CI | d_M | d_P | Transgress |
| --- | --- | --- | --- | --- | --- | --- |
| ALH | Mean | -0.03 | [-0.77, 0.70] | 0.80 | -0.67 | false |
| ALH | Skewness | 0.75 | [-0.02, 1.50] | -0.32 | 0.27 | false |
| ALH | Variance | 0.10 | [-0.64, 0.83] | 0.72 | -0.61 | false |
| **BCF** | **Mean** | **1.48** | **[0.65, 2.30]** | **0.45** | **-0.38** | **true** |
| BCF | Skewness | -0.39 | [-1.13, 0.36] | -0.69 | 0.59 | false |
| **BCF** | **Variance** | **2.30** | **[1.36, 3.21]** | **0.30** | **-0.26** | **true** |
| Head.Area | Mean | 0.16 | [-0.60, 0.91] | -0.52 | 0.47 | false |
| Head.Area | Skewness | -0.39 | [-1.15, 0.37] | 0.40 | -0.36 | false |
| **Head.Area** | **Variance** | **-1.41** | **[-2.24, -0.57]** | **0.24** | **-0.22** | **true** |
| **LIN** | **Mean** | **2.30** | **[1.36, 3.21]** | **-0.18** | **0.15** | **true** |
| **LIN** | **Skewness** | **-1.79** | **[-2.64, -0.93]** | **0.08** | **-0.07** | **true** |
| **LIN** | **Variance** | **2.03** | **[1.13, 2.91]** | **-0.37** | **0.32** | **true** |
| Midpiece.Area | Mean | -0.06 | [-0.81, 0.70] | -0.82 | 0.75 | false |
| Midpiece.Area | Skewness | -0.32 | [-1.07, 0.44] | 0.65 | -0.59 | false |
| Midpiece.Area | Variance | -0.06 | [-0.81, 0.69] | -0.50 | 0.45 | false |
| **STR** | **Mean** | **2.20** | **[1.28, 3.10]** | **-0.17** | **0.15** | **true** |
| **STR** | **Skewness** | **-1.96** | **[-2.83, -1.07]** | **0.21** | **-0.18** | **true** |
| STR | Variance | -0.55 | [-1.30, 0.20] | -0.51 | 0.43 | false |
| Tail.Length | Mean | 0.19 | [-0.56, 0.94] | -0.35 | 0.32 | false |
| Tail.Length | Skewness | 0.24 | [-0.52, 0.99] | 0.12 | -0.11 | false |
| Tail.Length | Variance | -0.60 | [-1.36, 0.17] | -0.76 | 0.69 | false |
| VAP | Mean | 1.16 | [0.36, 1.95] | 0.53 | -0.45 | false |
| VAP | Skewness | -0.92 | [-1.69, -0.14] | -0.47 | 0.40 | false |
| VAP | Variance | 1.06 | [0.27, 1.83] | 0.71 | -0.60 | false |
| VCL | Mean | 0.42 | [-0.33, 1.16] | 0.80 | -0.67 | false |
| VCL | Skewness | -0.35 | [-1.09, 0.39] | -0.65 | 0.55 | false |
| VCL | Variance | 0.06 | [-0.68, 0.80] | 0.82 | -0.70 | false |
| **VSL** | **Mean** | **1.94** | **[1.05, 2.81]** | **0.14** | **-0.12** | **true** |
| **VSL** | **Skewness** | **-1.29** | **[-2.08, -0.48]** | **-0.32** | **0.27** | **true** |
| **VSL** | **Variance** | **2.17** | **[1.25, 3.06]** | **0.07** | **-0.06** | **true** |
| **WOB** | **Mean** | **1.43** | **[0.60, 2.23]** | **-0.12** | **0.10** | **true** |
| WOB | Skewness | -0.48 | [-1.23, 0.27] | -0.06 | 0.05 | false |
| WOB | Variance | 0.68 | [-0.08, 1.43] | -0.32 | 0.27 | false |

**Table S4:** Pairwise post-hoc contrasts of integration intensity (V_rel_) across morphological and kinematic traits. Pairwise differences in integration magnitude were evaluated using linear models followed by estimated marginal means (emmeans). Contrasts, standard errors, and *P*-values were calculated for all species combinations, with *P*-values adjusted for multiple comparisons using Tukey’s HSD method. Contrasts were classified as statistically significant at an adjusted threshold of *P* < 0.05 (bolded). Together, these comparisons highlight how hybrids maintain comparable levels of morphological integration while showing kinematic integration that is elevated over non-hybrid species.

| Trait_Type | Comparison | Estimate | SE | p_value |
| --- | --- | --- | --- | --- |
| Morphology | generalist - molluscivore | -0.0048 | 0.0102 | 0.990 |
| Morphology | generalist - scale-eater x molluscivore | -0.0069 | 0.0102 | 0.960 |
| Morphology | generalist - scale-eater | -0.0089 | 0.0099 | 0.897 |
| Morphology | generalist - wide-mouth | -0.0197 | 0.0102 | 0.313 |
| Morphology | molluscivore - scale-eater x molluscivore | -0.0021 | 0.0106 | 1.000 |
| Morphology | molluscivore - scale-eater | -0.0041 | 0.0104 | 0.995 |
| Morphology | molluscivore - wide-mouth | -0.0149 | 0.0106 | 0.629 |
| Morphology | scale-eater x molluscivore - scale-eater | -0.0020 | 0.0104 | 1.000 |
| Morphology | scale-eater x molluscivore - wide-mouth | -0.0128 | 0.0106 | 0.750 |
| Morphology | scale-eater - wide-mouth | -0.0108 | 0.0104 | 0.835 |
| Kinematics | generalist - molluscivore | 0.0245 | 0.0179 | 0.650 |
| Kinematics | generalist - scale-eater x molluscivore | -0.0433 | 0.0184 | 0.143 |
| Kinematics | generalist - scale-eater | 0.0222 | 0.0172 | 0.698 |
| Kinematics | generalist - wide-mouth | -0.0018 | 0.0179 | 1.000 |
| **Kinematics** | **molluscivore - scale-eater x molluscivore** | **-0.0678** | **0.0187** | **0.006** |
| Kinematics | molluscivore - scale-eater | -0.0023 | 0.0176 | 1.000 |
| Kinematics | molluscivore - wide-mouth | -0.0263 | 0.0183 | 0.605 |
| **Kinematics** | **scale-eater x**<br>**molluscivore - scale-eater** | **0.0655** | **0.0180** | **0.006** |
| Kinematics | scale-eater x<br>molluscivore - wide-mouth | 0.0415 | 0.0187 | 0.191 |
| Kinematics | scale-eater - wide-mouth | -0.0240 | 0.0176 | 0.652 |

**Table S5:** Post-hoc pairwise PERMANOVA results testing divergence in individual kinematic correlation architecture across lineages. Pairwise PERMANOVA tests (pairwise.adonis) comparing male-level kinematic correlation matrices among purebred species (generalist [A], wide-mouth [S], molluscivore [M], scale-eater [P]) and hybrids (MxP). Male-level integration matrices across all kinematic traits were upper-triangle flattened and Fisher *z*-transformed to linearize correlations and satisfy normality assumptions. Multivariate Euclidean distances were then calculated across transformed correlation vectors. *P*-values were adjusted for multiple testing via Bonferroni correction and significant differences between groups (*P* < 0.05) were bolded.

| Pairs | Df | SS | F | R2 | p_raw | p_adj | sig |
| --- | --- | --- | --- | --- | --- | --- | --- |
| P vs M | 1 | 3.639 | 3.446 | 0.135 | 0.013 | 0.13 |  |
| P vs A | 1 | 0.733 | 0.717 | 0.030 | 0.554 | 1.00 |  |
| P vs S | 1 | 1.803 | 1.888 | 0.079 | 0.128 | 1.00 |  |
| **P vs MxP** | **1** | **8.855** | **9.451** | **0.310** | **0.001** | **0.01** | **\*** |
| M vs A | 1 | 3.095 | 2.792 | 0.117 | 0.027 | 0.27 |  |
| M vs S | 1 | 2.021 | 1.944 | 0.089 | 0.102 | 1.00 |  |
| **M vs MxP** | **1** | **16.050** | **15.672** | **0.452** | **0.001** | **0.01** | **\*** |
| A vs S | 1 | 1.516 | 1.512 | 0.067 | 0.188 | 1.00 |  |
| **A vs MxP** | **1** | **6.477** | **6.568** | **0.247** | **0.001** | **0.01** | **\*** |
| **S vs MxP** | **1** | **11.083** | **12.215** | **0.391** | **0.001** | **0.01** | **\*** |

